# Cortex-wide single-cell activity mapping in freely moving mice

**DOI:** 10.64898/2026.10.08.755518

**Authors:** Changliang Guo, Mian Xie, Guohua Qiu, Jian Lu, Yuzhen Liu, Liying Qu, Yuanyuan Huang, Zhenran Guo, Jing Tian, Jingren Lin, Baoyi Zhang, Xuefei Zhang, Junjie Zhao, Xiang Liu, Haochen Long, Jingyi Ma, Bo Zhou, Xianxin Ye, Pingping Zhao, Jingjie Cao, Zhengan Lin, Zhimiao Zhang, Weijie Deng, Jian Wang, Wei Wu, Junwen Wang, Xiaohong Peng, Shiqun Zhao, Shuai Na, Haoyu Li, Wenhao Liu, Qiang Fu, Zengcai V. Guo, Weisong Zhao, Liangyi Chen

**Affiliations:** National Biomedical Imaging Center, College of Future Technology, Peking University, Beijing, China; State Key Laboratory of Membrane Biology, Institute of Molecular Medicine, Peking-Tsinghua Center for Life Sciences, Beijing, China; IDG/McGovern Institute for Brain Research, Peking University, 100081, Beijing, China; School of life Sciences, Tsinghua University, Beijing, China; Academy for Advanced Interdisciplinary Studies, Peking University, Beijing, China; IDG/McGovern Institute for Brain Research, Tsinghua University, 100084, Beijing, China; School of Basic Medical Sciences, Tsinghua University, 100084, Beijing, China; Tsinghua-Peking Center for Life Sciences, 100084, Beijing, China; Innovation Photonics and Imaging Center, School of Instrumentation Science and Engineering, State Key Laboratory of Matter Behaviors in Space Environment, Frontier Science Center for Interaction between Space Environment and Matter, Harbin Institute of Technology, Harbin, China; Division of Applied Oral Sciences and Community Dental Care, Faculty of Dentistry, The University of Hong Kong; Beijing Institute of Collaborative Innovation, Beijing, China; Department of Neurology, David Geffen School of Medicine, University of California, Los Angeles, Los Angeles, CA, USA; Changchun Institute of Optics, Fine Mechanics and Physics, Chinese Academy of Sciences, Changchun 130033, China; University of Chinese Academy of Sciences, Beijing 100049, China; State Key Laboratory of Advanced Manufacturing for Optical Systems, Changchun 130033, China; The First Hospital of Jilin University, Changchun 130021, China; School of Basic Medical Sciences, Shenzhen University Medical School, Shenzhen 518060, China; PKU-Nanjing Institute of Translational Medicine, Nanjing, 211800, China; Academy for Advanced Interdisciplinary Studies, Peking University, Beijing, 100871, China.6.7; Beijing City Key Laboratory of Cardiometabolic Molecular Medicine, Peking University, Beijing,100871,China

**Author notes:** These authors contributed equally (Changliang Guo, Mian Xie, Guohua Qiu, Jian Lu, Yuzhen Liu).

## Abstract

How the cortex supports complex, naturalistic behavior can be revealed only in freely moving animals. Although head-mounted microscopes promise access to the neural dynamics underlying such behavior, fundamental trade-offs among field of view, spatial resolution, device weight, and background contamination have prevented cortex-wide single-neuron recording, leaving distributed cortical coordination largely inaccessible. To meet these challenges, we introduce **COMET (C**ortex-wide **O**bservational **M**iniature **E**pifluorescence **T**echnique), a 3.75-g head-mounted mesoscope that integrates an aberration-controlled wide-field optical design, uniform high-efficiency excitation and single-coaxial power-and-data transmission with an end-to-end denoising and signal-extraction pipeline. This enables cortex-wide calcium imaging at single-cell resolution in freely moving mice. COMET captures activity from >15,000 layer 2/3 neurons per 5-min recording over fields of view up to 10 mm in diameter while preserving natural behavior. Benchmarking against benchtop two-photon microscopy demonstrates high spatial concordance of detected neurons and matched event-level calcium dynamics. We apply COMET to social interaction to quantify region-specific engagement and to an auditory trace fear-conditioning task that requires natural movement and behavioral expression, revealing time-ordered changes in regional firing-rate structure and functional connectivity preceding a global conditioned response. COMET provides an open, lightweight hardware–software framework for linking single-neuron activity to cortex-scale network dynamics during freely expressed behaviors under changing internal or external conditions.

## Main

Neuronal computation in complex and naturalistic behavior emerges from distributed activity patterns that span multiple cortical areas. Capturing these dynamics at the level of individual neurons—rather than regional averages—requires recording from large neuronal populations simultaneously, ideally across the dorsal cortex. Although layer 2/3 is a key locus for intercortical communication and cortical output routing in canonical circuits^1,2^, existing experimental access to L2/3 dynamics remains constrained by the limited spatial coverage and/or mobility of available imaging approaches^3^.

Genetically encoded calcium indicators paired with fluorescence microscopy provide a scalable route to large-scale functional mapping^4,5,6,7^, and head-fixed preparations have enabled cellular-resolution recordings across multiple cortical regions^8–12^. However, immobilization precludes many ethologically relevant behaviours and alters sensorimotor loops, motivating head-mounted systems for freely moving animals. Miniature two-photon microscopes offer optical sectioning and high resolution in motion^5,13–21^, but their field of view is fundamentally limited by scanning throughput and photon efficiency, restricting cortex-scale coverage. One-photon miniscopes can expand field of view^20^, yet achieving **whole-cortex, single-cell resolution** in a device light enough for unrestrained use has remained difficult because wide-field epifluorescence amplifies background and demands stringent optical aberration control, efficient light collection, lightweight mechanics, and analysis methods that can robustly separate neurons under low SBR conditions.

Here we introduce **COMET** (**C**ortex-wide **O**bservational **M**iniature **E**pifluorescence **T**echnique), a **3.75 g** head-mounted mesoscope that closes this gap by co-designing **wide-field, aberration-controlled optics**, a compact mechanical/electronics architecture, and an end-to-end computational pipeline for artifact removal, denoising, neuron extraction and atlas registration. COMET enables full-frame imaging over fields up to **10 mm** in diameter at **9.3 Hz** and resolves activity from **>15,000 neurons** across most of the dorsal cortex in freely moving mice, while minimizing cable burden through a single-coax power-and-data link. Using this platform, we map cortex-wide single-cell dynamics during spontaneous social interaction to quantify region-specific engagement at single-cell resolution and an auditory trace fear-conditioning paradigm to reveal temporally ordered changes in regional activity and functional connectivity during learning. Together, COMET provides a lightweight, scalable hardware–software framework for linking single-neuron activity to cortex-scale network organization during naturalistic behavior.

## Results

### COMET and computational pipeline enable cortex-wide imaging with single-cell resolution

To enable cortex-wide calcium imaging in freely behaving mice, a head-mounted microscope must satisfy three coupled requirements: (1) a lightweight, compact form factor (<4 g) that does not measurably perturb natural behavior; (2) single-cell spatial resolution (⩽10 µm) sufficient to resolve individual neuronal activity; and (3) a large field of view (⩾ 8 mm diameter) spanning most of the dorsal cortex. Meeting these specifications required co-optimization of four elements: an aberration-controlled wide-field optical train, lightweight yet rigid mechanics, low-noise high-throughput electronics, and computational processing that can reliably extract single-neuron signals from high-background epifluorescence data.

COMET integrates three miniaturized hardware modules—an optics module, a baseplate/illumination module, and a custom rigid–flex PCB module (**Fig. 1a; Supplementary Video 1**). Uniform excitation across the full field is achieved using eight LEDs (LXZ1-PB01, LUXEON) coupled to stacked excitation filters (ET470/40x, Chroma; non-coated sides facing) and a diffuser layer to homogenize illumination (**Figure 1b**). We designed a five-lens miniature objective (object-space NA = 0.083) that supports fields up to 10 mm in diameter (78.54 mm²) with near diffraction-limited performance in simulation (**Extended Data Fig. 1**). In contrast, a more compact four-lens alternative exhibited enlarged off-axis point spread functions (PSFs) at 4 mm field radius and a faster drop in modulation transfer at higher spatial frequencies (**Extended Data Fig. 1a**). An electrowetting lens (EWL; A-25H, Varioptics/Corning) positioned near the pupil plane provides electronic focus adjustment (**Extended Data Fig. 1a left**). At 0.468× magnification, optical distortion and aberrations remain low across the field, as confirmed using a 500-µm grid target (**Extended Data Fig. 1b**).

**Fig. 1.**
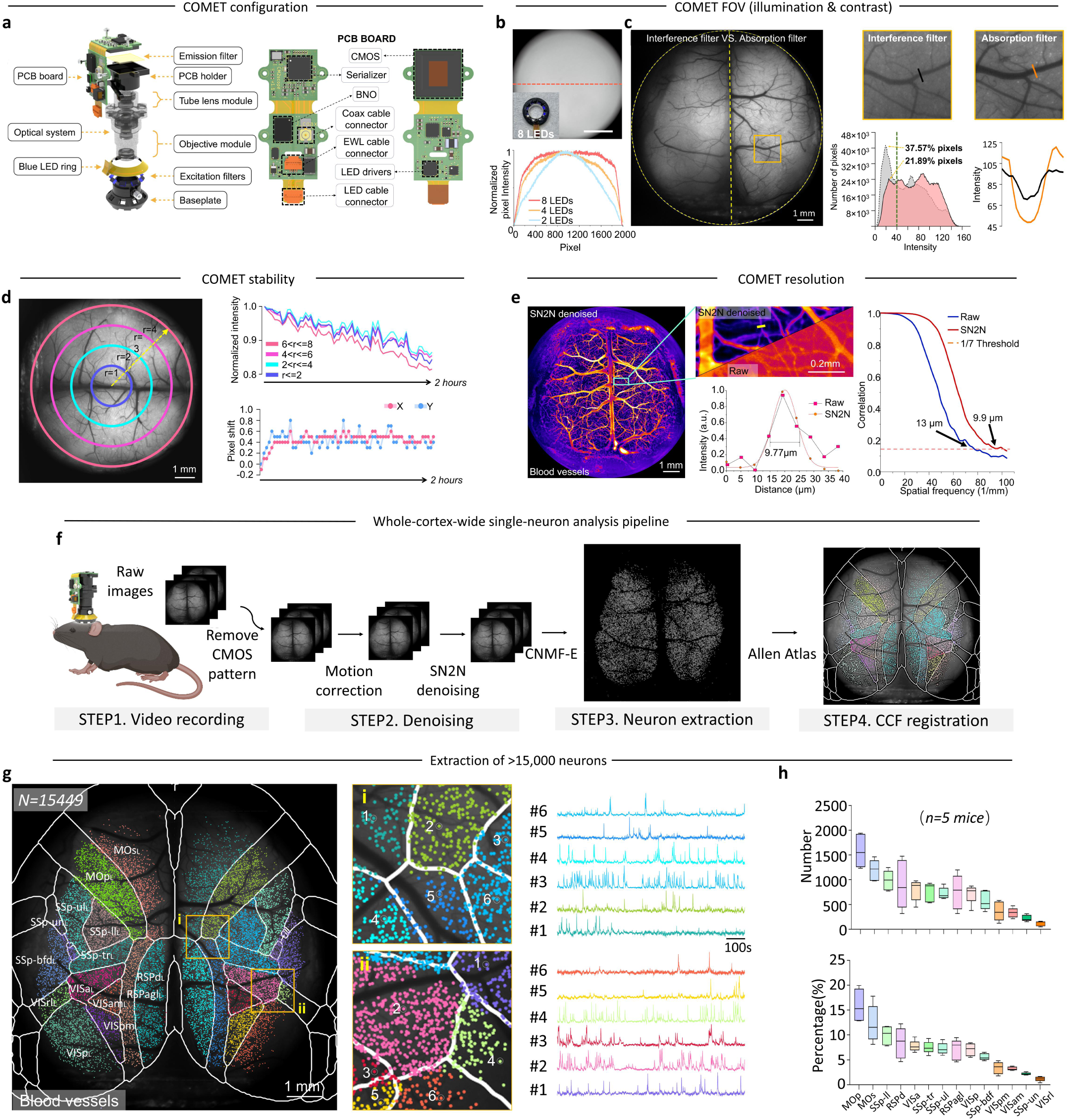
COMET achieves cortex-wide imaging with single-cell resolution across up to a 10-mm FOV in diameter. **a**, System overview. Left: Exploded view schematic of the COMET, highlighting three core modular components: (i) Electronic control unit: Flexible PCB board integrating a blue LED excitation ring; (ii) Optical imaging module: Objective lens assembly (NA 0.083), 4-element tube lens system (0.468× magnification), bandpass excitation filter, angle-insensitive emission filter, and (iii) Mechanical housing: 3D-printed baseplate and PCB holder (black resin, Formlabs), and magnesium-aluminum alloy structural components for weight optimization. Right: Rigid-flex PCB design comprising (i) a control electronics board including two LED drivers with I²C-controlled digital potentiometer for 8 excitation LEDs, EWL driver, BNO055 orientation sensor, and high-speed serializer for coaxial data transmission, and (ii) an imaging module with 5-MP CMOS sensor (MT9P031). Scale bar: 1 cm. **b,** Uniform illumination system. Eight LEDs coupled with dual bandpass filters (16 total) and paper diffusers achieve homogeneous excitation across the full FOV. Scale bar: 2 mm. **c,** Emission filtering strategy. Quantitative comparison shows absorption filters provide superior contrast (21.89% vs 37.57% pixels <40 grayscale) over interference filters by eliminating angle-dependent wavelength shifts. Blood vessel visibility demonstrates enhanced contrast (insets). Scale bar: 1 mm. **d,** COMET enables at least a 2-hour recording with only slight bleaching and less than 1 pixel shift of the recordings.The first frame from a 2-hour recording of a freely moving mouse, circled with different radius from 1 to 4 mm in the FOV (**left**). Normalized intensity change in the FOV, showing a less 20% bleaching within a 2-hour recording. Pixel shift is less than 1 pixel both in x and y directions within 2-hour recording (**right**). **e,** COMET enables blood vessel imaging in the cortex of freely moving mice following FITC–dextran injection. Left: Fluorescence image of cortical blood vessels after denoising using the SN2N algorithm (**left**); Middle: The zoomed-in region highlights the comparison before and after SN2N processing (**top**). An intensity profile across a representative blood vessel and the corresponding Gaussian fit indicate an achievable full width at half maximum (FWHM) of 9.77 µm (**bottom**). Right: Fourier ring correlation (FRC) analysis of the denoised images. **f**, Processing workflow. (1) Raw data acquisition; (2) CMOS pattern noise removal (died pixel correction); (3) Motion artifact compensation; (4) Denoising using SN2N algorithm; (5) Neuron extraction via CaImAn (CNMF-E); (6) CCF registration to Allen Reference Atlas. **g,** CCF-registered neurons (n=15,449, 3000 frames for 5-min recording) from one mouse, with insets showing: (i) motor/somatosensory ROI (MOs, MOp, SSP-ll/tr, RSPagl/d) and (ii) visuoparietal ROI (SSP-tr/bfd, VISa/rl/p/am/pm). Normalized activity traces from 6 representative neurons per ROI shown (right). Scale bar: 1 mm. **h,** Cortical distribution of neuron counts (n=5 mice), demonstrating predominant localization to central regions (MOs/MOp) with fewer cells at FOV periphery (SSP-un, VISrl).

To suppress excitation leakage and backscatter—especially problematic in wide-field epifluorescence—we used an angle-insensitive absorption emission filter (Wratten dye filter; 10 × 10 × 0.1 mm, Kodak Wratten #54-467, Edmund Optics) in place of an interference filter. Unlike interference filters whose passband shifts at high incidence angles, the absorption filter maintains rejection across angularly distributed stray light, improving image contrast and homogeneity across the field (**Fig. 1c**), enabling stably at least 2-hour recordings with slight bleaching and less than 1 pixel shift. (**Fig. 1d**). We assessed effective in vivo resolution by imaging cortical vasculature in freely moving mice after intravenous FITC–dextran injection. After denoising with SN2N^22^, COMET resolved vessels as small as 9.77 µm in diameter, and Fourier ring correlation independently yielded an effective resolution of 9.9 µm (**Fig. 1e**). Across operating modes, resolution is determined by objective NA and sampling: USAF target measurements showed 4.39 µm at 3.96 mm FOV, 6.21 µm at 5.95 mm FOV, and 8.77 µm at 8.94 mm FOV (**Extended Data Fig. 1b, middle**). Optical magnification decreased with working distance from 1.3× (0 mm) to 0.45× (6.5 mm) (**Extended Data Fig. 1c, 1d**). Consistently, 2-µm bead imaging yielded average bead FWHM values of 7.21 µm at the field center, 7.64 µm at 2 mm off-axis, and 9.79 µm at 4 mm off-axis (**Extended Data Fig. 1e**).

Mechanical and electronic subsystems were engineered for rigidity at minimal mass. The optical housing was machined from magnesium–aluminium alloy (AZ31B; 1.77 g cm⁻³) and electrochemically blackened to suppress internal reflections and scattered light. The baseplate was 3D-printed in light-absorbing black resin (Formlabs Form 3; 25-µm layers) to (i) precisely position the LED/filter illumination ring, (ii) provide a low-profile cemented interface to the skull, and (iii) stabilize the imaging module during complex behaviors (**Fig. 1a, Extended Data Fig. 2**). The rigid–flex PCB comprises three functional subcircuits: a control board for EWL tuning, inertial sensing (BNO055, Bosch), and power-over-coax plus serialization; an imaging board hosting a 5-MP monochrome CMOS sensor (MT9P031, Onsemi); and a flexible excitation board driving eight independently addressable LEDs. These are linked by a high-density embedded flex interconnect, enabling robust high-bandwidth transmission with mechanical compliance (**Fig. 1a, Extended Data Fig. 2**). The PCB is mounted on custom 3D-printed shells with M1 stainless screws to maintain alignment and electrical isolation while minimizing mass (**Extended Data Fig. 2**).

To reduce cabling burden and behavioral interference, COMET delivers power, bidirectional control (I²C), and high-bandwidth image data through a single 50-Ω coaxial cable (CW2040-3650SR, Cooner Wire) using power-over-coax filtering and a SerDes link (**Fig. 1a; Extended Data Fig. 2a,2b**). The system interfaces with the UCLA open-source Miniscope DAQ software, enabling real-time control of excitation intensity, EWL focus (via MAX14515 driver, Maxim), and sensor parameters (including up to 8× analog gain), as well as synchronization with behavioural cameras and external triggers^16,23^. COMET records full-frame video at 9.3 Hz (2592 × 1944) and can reach ∼30 Hz via ROI cropping or binning, balancing cortex-scale coverage with single-cell spatial resolution for freely moving calcium imaging. At a total weight <4 g, COMET’s lightweight and stable design enable natural behavior (**Extended Data Fig. 3a,3b**). COMET also allows stably cross-day recordings, validated by tracking cortical cells for 3 continuous days. (**Extended Data Fig. 3c**).

Multiphoton imaging typically yields high signal-to-background ratios (SBRs) because nonlinear excitation provides intrinsic optical sectioning^14,15,17,18,24^. In contrast, single-photon epifluorescence integrates fluorescence from both in-focus and out-of-focus planes, reducing SBR and complicating neuron identification and trace extraction. To overcome this limitation, we established an end-to-end computational pipeline optimized for COMET recordings from transgenic mice with sparse GCaMP6f expression in layer 2/3 neurons^25^ (**Methods**, **Fig. 1f, Supplementary Video 3**). The workflow comprises four steps: (1) acquisition and sensor artifact correction, including removal of CMOS pattern noise associated with heat-induced dead pixels, (2) motion correction and denoising using SN2N^22^ to improve effective SBR, (3) neuron extraction using a CNMF-E–based approach^26,27^ to obtain spatial footprints and calcium traces, and (4) atlas registration to the Allen Mouse Common Coordinate Framework (CCF)^28^ to assign extracted neurons to anatomical regions. This pipeline enables scalable single-neuron extraction and anatomical referencing for cortex-wide, region-resolved analyses of neuronal population dynamics during behavior, and transforms raw wide-field recordings into spatially organized single-neuron activity maps suitable for cross-animal and cross-study comparisons.

Finally, we registered COMET-extracted neurons to the Allen CCF to enable region-resolved analyses. In one representative dataset, 15,449 neurons were assigned to 28 cortical regions (**Fig. 1g, left**). We then defined two representative multi-region ROIs—ROI-i (MOs, MOp, SSP-ll, SSP-tr, RSPagl, RSPd) and ROI-ii (SSP-tr, SSP-bfd, VISa, VISrl, VISp, VISam, VISpm) (**Fig. 1g, middle**)—and extracted example traces from randomly selected neurons in each ROI to illustrate comparably high SNR across widely distributed cortical areas (**Fig. 1g, right**). Neuron counts were highest in centrally located regions such as MOs and MOp, with fewer neurons detected at the periphery of the field (**Fig. 1h**).

### Validation of high-fidelity single-neuron extraction from COMET against two-photon microscopy

The denoising workflow substantially improved visual quality and single-neuron separability relative to unprocessed data (**Fig. 2a**, **Fig. 2b-left**). Using the temporally averaged image from each 5-min recording (∼3,000 frames) as a reference, we quantified these improvements and observed significant increases in both structural similarity (SSIM) and peak signal-to-noise ratio (PSNR) after denoising (**Fig. 2b, middle and right**). To benchmark the fidelity of neuron extraction from COMET recordings, we imaged the same cortical regions in the same mice using a commercial two-photon (2P) microscope (N10XW-PF-10X Nikon CFI Plan Fluorite Objective, 0.30 NA, 3.5 mm WD, FVMPE-RS, Olympus) (**Fig. 2c, Extended Data Fig. 4, Supplementary Video 4**). In head-fixed preparations, we acquired an 11-plane 2P stack spanning depths of 20–170 μm in 15-μm steps. After SN2N-based denoising^22^, 2P data yielded 3,659 neurons, while COMET detected 1,870 neurons in the corresponding regions (n = 2 mice). Across datasets, an average of 87.17% of COMET-identified neurons colocalized with neurons resolved by 2P imaging (**Extended Data Fig. 4, Fig. 2d, bottom right**). Matched neurons exhibited highly concordant spontaneous calcium dynamics across modalities(**Fig. 2d, left**), and neither ΔF/F amplitudes (n = 20 neurons) nor event rates differed significantly between COMET and 2P recordings (n.s., Mann–Whitney test; **Fig. 2d, top right and middle right**), supporting the reliability of COMET for single-cell calcium signal detection.

**Fig. 2.**
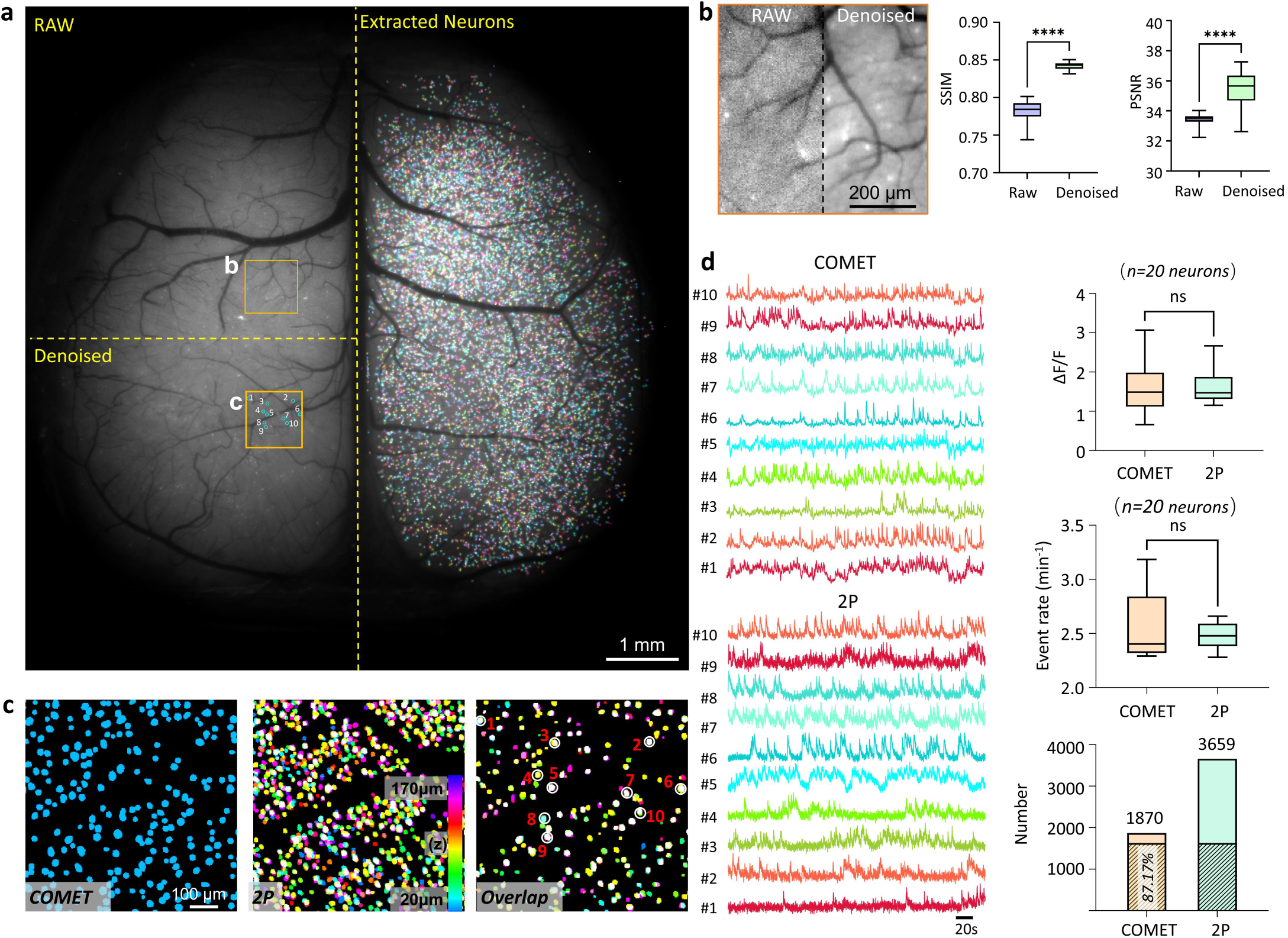
COMET’s computational pipeline enables high-fidelity neuronal signals extraction. **a**, Representative 5-min recording from a mouse showing: raw subregion (top left), denoised output (bottom left), and extracted neuron ROIs (right). Scale bar: 1 mm. **b**, Quantitative denoising performance. Zoomed region (orange box in **a**) comparing raw vs. processed images with corresponding SSIM (a mean value 0.782 raw →a mean value 0.842 denoised) and PSNR (33.437 raw → 35.512 denoised) improvements (middle/right). ****p<0.001, paired t-test. **c,** Validation against two-photon microscopy. Neuron footprints from COMET (left), 11-plane two-photon stack (middle), and overlap (right; 87.1% concordance). Scale bar: 100 µm. **d,** Functional validation. Normalized calcium traces from 10 matched neurons demonstrate highly concordant temporal dynamics between COMET (top left) and two-photon microscopy (bottom left). Quantitative population analysis revealed no significant differences in either ΔF/F amplitudes (median 1.488 COMET → median 1.473 2P, p>0.05, Mann-Whitney test) or event rates (a median value 2.403 COMET → a median value 2.480 2P, p>0.05, Mann-Whitney test) between modalities (top/middle right). Spatial mapping confirmed 87.1% correspondence in detected neuronal positions (bottom right), validating COMET’s accuracy for single-cell resolution imaging.

### Social interaction paradigm using COMET

Social behavior recruits distributed circuits to integrate sensory cues with internal state and to coordinate motor output ^30^. To demonstrate COMET’s ability to capture cortex-wide single-cell activity during naturalistic behavior, we imaged layer 2/3 (L2/3) activity in transgenic mice sparsely expressing GCaMP6f^25^ during a simple two-animal social interaction assay (**Methods**). A COMET-mounted mouse explored an open-field arena for 5 min, after which a second mouse was introduced and interactions were recorded for an additional 5 min (**Fig. 3a**). Across five mouse pairs, COMET captured high-SNR activity from >10,000 neurons distributed across 26 cortical region (**Fig. 3b,3c**). Social interaction epochs were manually annotated from overhead behavioral video (**Methods**).

**Fig. 3.**
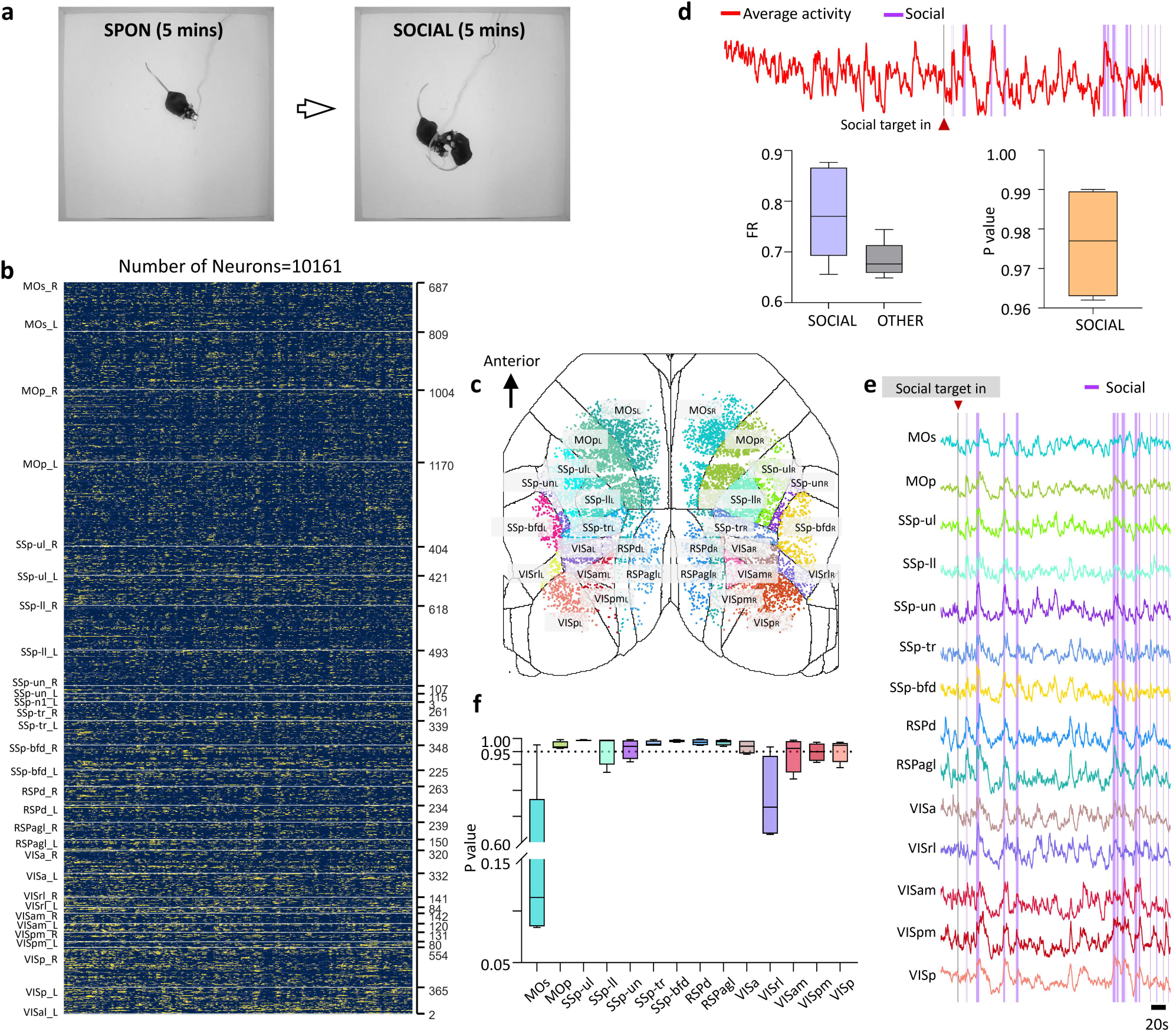
Global cortical synchronization during social interaction. **a,** Behavioral paradigm: 5-minute free exploration and 5-miniute social interaction between mice. **b,** Normalized calcium activity (ΔF/F) of 10,161 neurons across 30 cortical regions. **c,** Spatial organization: Neuron footprints of CCF-registered neurons. **d,** Analysis with population-averaged activity of individual neurons reveals elevated firing rates (FR) during social interaction (left) compared to baseline (middle), with consistent effects across all mice (permutation test, right). **e,** Normalized region-specific population-averaged activity demonstrates distributed engagement. **f,** P-values of six regions (MOp, SSP-ul, SSP-tr, SSP-bfd, RSPd, RSPagl) show particularly high involvement (p < 0.05, permutation test).

We first asked whether global cortical activity increased during social interaction (**Fig. 3d, top**). Event-derived firing rates (FR; **Methods**) were higher during social than non-social periods (0.7777 ± 0.09113 vs. 0.6842 ± 0.03587; **Fig. 3d, bottom left**). This effect was supported by permutation testing, in which the observed difference exceeded the majority of shuffled surrogates across all five animals (empirical significance score > 0.95 across all five mice; **Fig. 3d, bottom right**). We then assessed regional engagement using an activity-based significance score per region (**Fig. 3e-3f**). Six regions—MOP, SSP-ul, SSP-tr, SSP-bfd, RSPd, and RSPagl—consistently exhibited high engagement (p > 0.95, **Fig. 3f**), consistent with contributions from somatosensory cortex to tactile/social cues^31^ and retrosplenial cortex to social-state regulation^32^.

### Revealing cross-scale memory formation during trace fear conditioning with COMET

Trace fear conditioning (TFC) probes how the brain links temporally separated events, as animals learn to associate a tone (CS) with a delayed footshock (US) while expressing spontaneous behavioral responses such as freezing, a conditioned response that is most robustly observed in freely moving preparations. While head-fixed setups can facilitate precise circuit measurements, they impose constraints on naturalistic movement and behavioral expression that can alter stimulus – response dynamics and underlying neural activity. Although hippocampal mechanisms bridging the trace interval are well studied^33–35^, how distributed cortical activity and inter-regional coupling evolve across learning trials in freely moving animals remains unclear. We therefore performed COMET imaging during TFC in freely moving mice with L2/3-specific GCaMP6f expression (n = 5; **Methods**). After 240 s of spontaneous activity (SPON), mice underwent five CS–US trials (20-s tone, 18-s trace, 2-s shock), each followed by a 60-s post-shock period (L) (**Fig. 4a, 4b**; **Supplementary Video 5**).

**Fig. 4.**
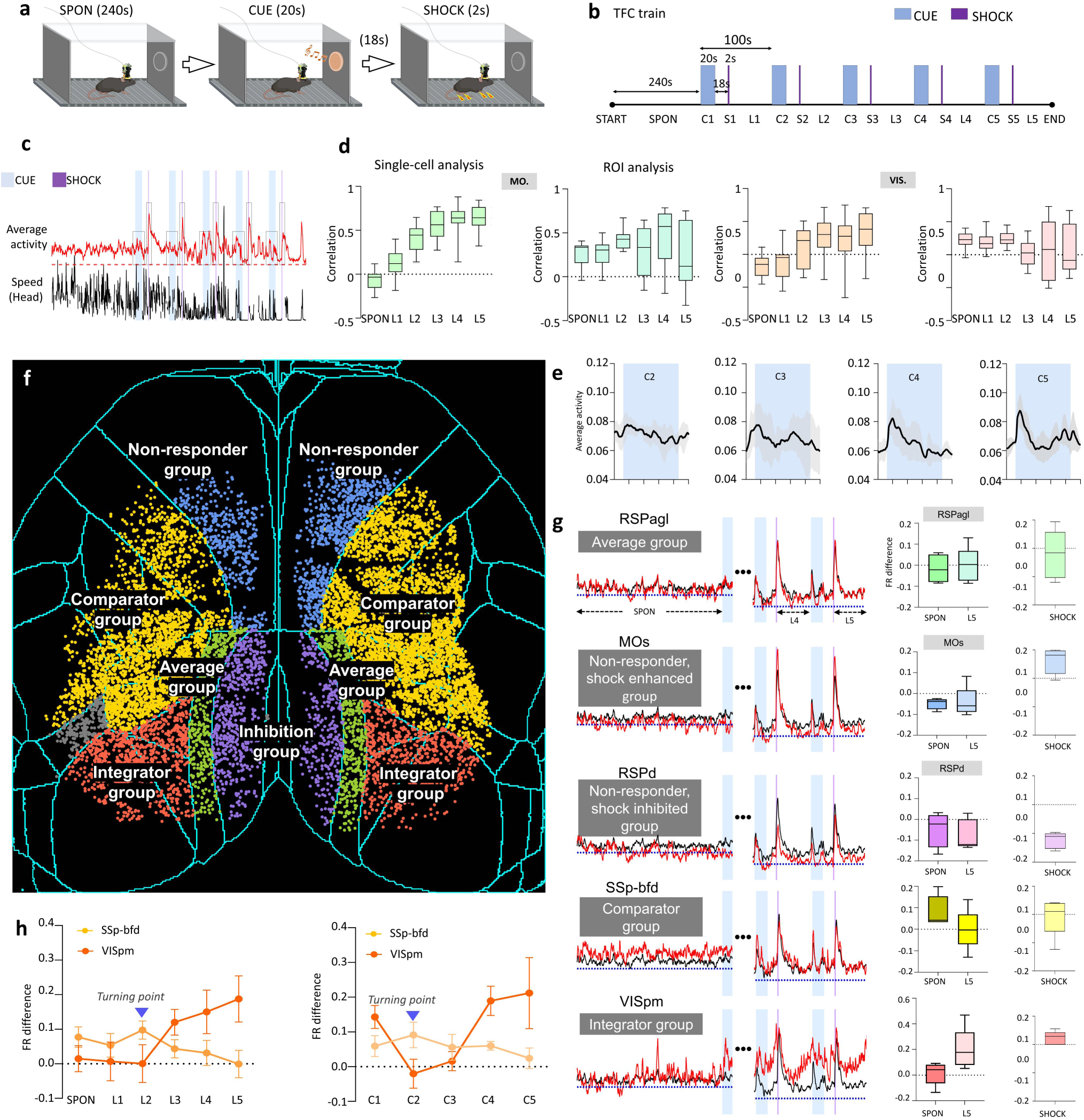
Functional heterogeneity in cortical activity during TFC. **a,** TFC paradigm structure. 20-s tone (C), 18-s trace interval, 2-s footshock (S), and 60-s post-shock period (L). **b,** Behavioral protocol showing 5 consecutive TFC trials terminated by a light signal. **c,** Neural-behavioral dynamics in Mouse #94. Population-averaged calcium activity (n=9,570 neurons; ΔF/F normalized) (**top**), concurrent running speed (head-movement derived) (**bottom**). **d,** Running-activity correlation analysis. Single-cell resolution Pearson correlation (r) during spontaneous (SPON) and post-shock (L1-L5) periods (**1^st^ and 3^rd^ column**) and ROI-based correlation (non-single-cell) for comparison (**2^nd^ and 4^th^ column**) in MO. and VIS. Cortical regions. **e,** Population dynamics during TFC. Population-averaged activity across all neurons (n = 5 mice), with enlarged view of C2–C5 trials showing increased response to conditioned stimulus (CS) onset (first 8 s of CUE). **f,** Cortical regions were categorized into five functional groups based on deviation from Population-averaged activity of whole-cortex: Average group (green colored) represented by RSPagl, Non-responder group (blue colored) represented by MOs, Inhibitory group (purple colored) represented by RSPd, Comparator group (yellow colored) represented by SSp-bfd, and Integrator group (red colored) represented by VISpm. **g,** Temporal profiles reveal group-specific modulation patterns during training compared to population-averaged activity of whole-cortex: Average group racks whole-cortex activity (**Row 1**); Non-responder group show stable baseline but elevated shock (US) response (**Row 2**); Inhibitory group remains suppressed below mean, including during US (**Row 3**); Comparator group exhibits declining divergence (**Row 4**); Integrator increase divergence post-CS (**Row 5**). **h,** Memory consolidation signature: A turning point in firing rate (FR) differences during L2/C2 may suggest plasticity onset.

Across regions, shocks induced transient cortex-wide activity increases (**Extended Data Fig. 5a, 5b, Fig. 4c**) that covaried with running speed estimated from head position (**Methods**). This coupling was region dependent: motor/somatosensory and retrosplenial regions showed strong and progressively increasing activity–speed correlations during post-shock periods (Pearson r peaking >0.5), whereas visual regions showed only modest changes (**Fig. 4d, Extended Data Fig. 5c, left column**). When we spatially smoothed COMET data to emulate lower-resolution, non-single-cell readouts used in prior mesoscale approaches^36^ (**Methods**), these temporally structured correlation changes were no longer detectable (**Fig. 4d, second and last columns, Extended Data Fig. 5c, right column**), underscoring the sensitivity gained by cortex-wide single-neuron measurements.

**Fig. 5.**
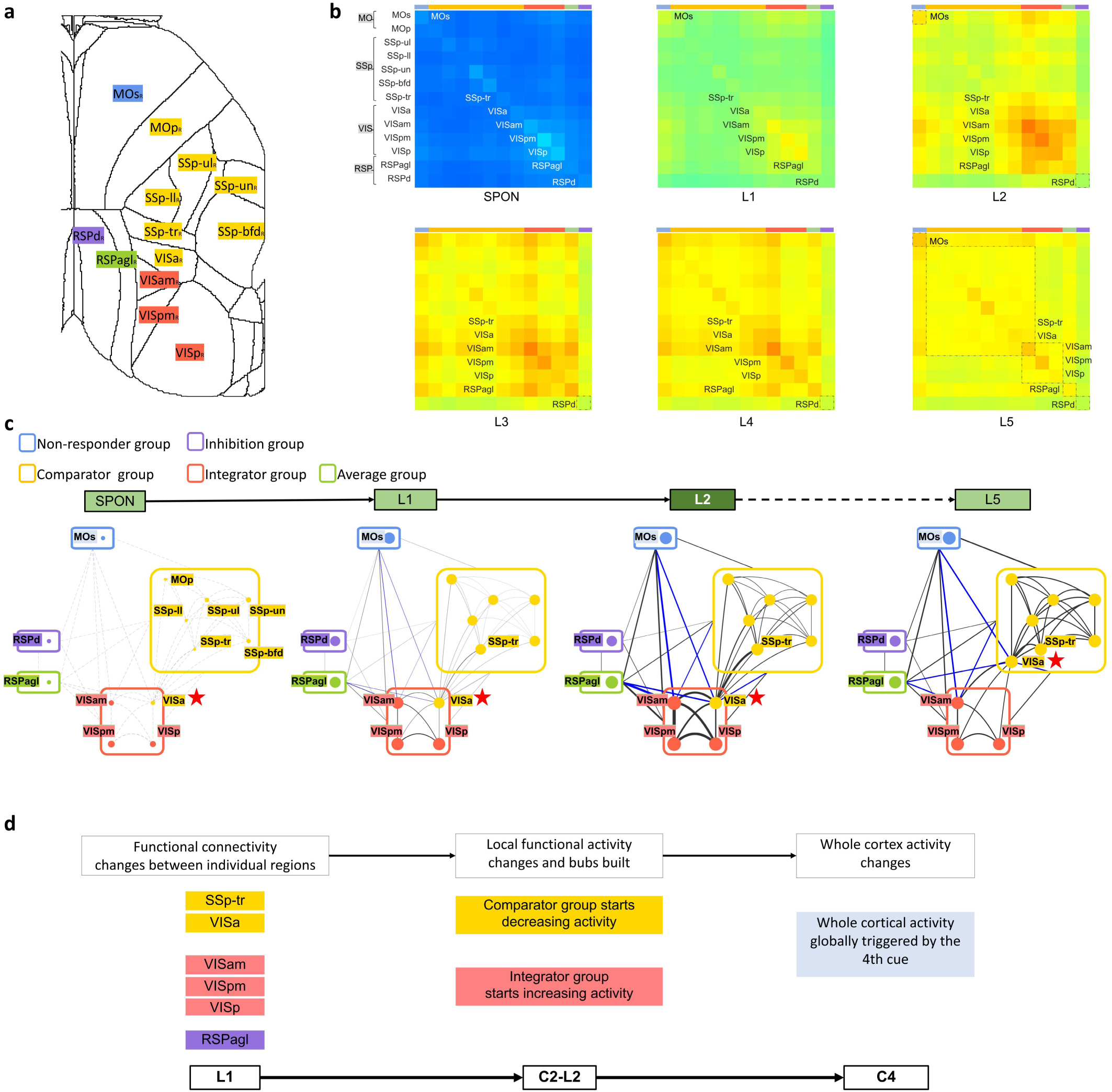
Network reorganization during TFC. **a,** Spatial distributions of 13 cortical regions (MOs, MOp, SSP-ul, SSP-ll, SSP-un, SSP-bfd, SSP-tr, VISa, VISam, VISpm, VISp, RSPagl, RSPd). **b-c,** Dynamic connectivity changes: During SPON, weak intra-regional correlations exist within visual (VIS), somatosensory (SSp-un/bfd), and motor (MO) areas. L1 initiates widespread intra-regional connectivity boosts and VISa’s functional reassignment. L2 strengthens long-range VISpm/am/p connections with non-adjacent RSPagl/MOs (represented by blue lines), while VISa couples with neighboring SSp-str. By L3-L5, visual network segregation stabilizes as comparator group (SSp) connectivity grows and VISam and RSPagl potentially serves as a hub gradually. **d,** Local synchronization is built after the first shock, advancing to cross-regional integration and coordinated activity changes among groups of brain regions after the second shock, and culminating in global cortical responses to the fourth cue.

We next tracked cortex-wide responses to the CS across training. While population-averaged CS-epoch activity increased with continued conditioning, clear temporal alignment to CS onset emerged only after the fourth trial (C4) (**Fig. 4e**). Using the population-averaged whole-cortex activity trace as a reference (REF), we compared regional dynamics to REF and grouped regions into five functional categories based on their deviation from REF during resting conditions and how this deviation evolved across trials (**Fig. 4e-4g)**. The average group (e.g., RSPagl) tracked REF throughout the session (**Fig. 4g, Row 1**). The shock-enhanced non-responder group (e.g., MOs) exhibited a selective increase during shocks (**Row 2**), whereas the shock-inhibited non-responder group (e.g., RSPd) remained suppressed, including during shocks (**Row 3**). Two groups showed time-dependent convergence toward REF: the comparator group (e.g., SSp-bfd) reduced its deviation beginning at L2 and plateaued by L4 (**Row 4**; **Fig. 4h, left**), whereas the integrator group (e.g., VISpm) showed the opposite trend (**Row 5**). VISpm further exhibited high activity/FR relative to REF at C1 that was transiently suppressed at C2–C3 (**Fig. 4h**, **right**), suggesting that the C2/L2 transition marks an early inflection point in distributed cortical dynamics during CS–US association.

Finally, we asked whether inter-regional coupling reorganized across training. We computed pairwise correlations between calcium activity traces of individual neurons within and across regions to construct weighted functional networks for each mouse and summarized network organization using mean degree centrality within and between regions (**Fig. 5a–5c**). Following the first shock (L1), coupling strengthened between VISa and SSp-tr (**Fig. 5c, L1**), and connectivity increased within the integrator-associated set and between VISam and regions in the average and shock-enhanced groups at L2 (**Fig. 5b,5c**). Because global arousal and movement can inflate correlation structure, we interpret these networks as proxies for functional coupling and focus on within-animal, epoch-to-epoch changes and region-specific patterns rather than absolute correlation magnitude. Notably, VISa was assigned to the comparator group by firing-rate deviation but displayed an integrator-like connectivity profile before L5 (**Fig. 5b,5c**), indicating that network-level reorganization can reveal relationships not apparent from mean activity alone. Overall, TFC dynamics progressed from local synchronization after the first shock to broader cross-regional integration after the second shock, culminating in a cortex-wide conditioned response by the fourth cue (**Fig. 5d**).

## Discussion

COMET expands the operating regime of head-mounted microscopy by enabling cortex-wide, single-cell calcium imaging in freely moving mice while avoiding the traditional trade-offs among field of view, resolution and weight that constrain existing miniature systems (**Table 1**)^10,16,20^. Crucially, this operating regime is required for studying complex, naturalistic behaviors, in which coordinated activity across multiple cortical areas unfolds in space and time and cannot be inferred from a limited subset of regions. This capability arises from the co-design of wide-field, aberration-controlled optics, lightweight mechanical integration, streamlined tethering, and an analysis pipeline explicitly tailored for high-background epifluorescence recordings. Together, these elements enable stable recordings from more than 15,000 neurons across fields of view up to 10 mm in diameter without disrupting natural behavior, providing a practical framework for mapping cortex-scale population dynamics at cellular resolution during complex behavior.

**Table 1.** Comparison of COMET to other miniaturized microscopes.

| Modalities | Weight<br>(g) | FOV<br>(mm) | Optical<br>resolution<br>($\mu\text{m}$) | Magnification | Single-cell<br>resolution | Needs<br>fibers |
| --- | --- | --- | --- | --- | --- | --- |
| COMET (Ours) | 3.75 | up to $\varnothing 10$ | 3.13<br>(effective<br>7–10) | 0.468× | Yes | No |
| SOMM<sup>49</sup> | 2.5 | $3.6 \times 3.6$ | 4 | $\sim 1\times$ | Yes | No |
| MinimScope<sup>36</sup> | 3.8 | $8 \times 10$ | 78 to 112 | 0.36-0.45× | No | No |
| cScope<sup>50</sup> | 35 | $7.8 \times 4$ | 14 | 0.7× | No | No |
| Kiloscope<sup>21</sup> | 1.4 | $4.8 \times 3.6$ | $\sim 10$ | $\sim 1\times$ | Yes | Yes |
| MiniXL<sup>20</sup> | 3.5 | $3.5 \varnothing$ | 4.4 | 1.1× | Yes | No |
| MiniLFOV<sup>16</sup> | 13.9 | $3.1 \times 2.3$ | 2.5 | 1.86× | Yes | No |
| | | $3.6 \times 2.7$ | 3.5 | 1.56× | | |

As a proof of capability, we first applied COMET to spontaneous social interaction, an ethologically relevant behavior that inherently requires unconstrained movement and engages distributed cortical processing. By quantifying region-specific engagement using an activity-based significance score, COMET enables a cortex-wide view of how natural social behavior recruits distinct cortical areas at single-cell resolution — an operating regime that has remained largely inaccessible to existing head-mounted imaging approaches. We further applied COMET to auditory trace fear conditioning and observed a staged emergence of cortex-wide responses across learning trials. Although fear memory formation is classically associated with hippocampal and amygdalar circuits^33–35^, we detected a pronounced global cortical response to the conditioned cue that emerged after repeated pairings, consistent with the ultra-dense interareal cortical network^37^ and reports that memory representations can be distributed across brain regions^38^. Using the whole-cortex response as a reference, we further identified regional groups whose activity evolved in distinct, time-ordered patterns relative to the global signal. In particular, a “comparator” group showed early elevations that diminished after L2, whereas an “integrator” group displayed transient suppression around C2–C3 followed by later recruitment. These observations support a model in which learning engages multi-stage, cross-scale cortical recruitment, progressing from local processing to broader integration of cue-related representations.

Single-cell resolution was essential for resolving these dynamics and for interrogating network organization beyond mean activity levels. Correlation-based functional networks constructed from individual neuronal traces revealed region-specific reconfiguration that was not apparent from averaged calcium signals alone. For example, VISa exhibited an early shift in its connectivity profile following the first shock, marking one of the earliest detectable network-level transitions despite minimal changes in its mean activity or firing rate. Similarly, RSPagl and MOs showed connectivity changes without obvious deviations from the global mean at L1. Notably, RSPagl within retrosplenial cortex—a region with extensive visual and hippocampal connectivity^39–44^—emerged as a candidate hub for cue–context integration, consistent with prior evidence implicating retrosplenial cortex in trace fear extinction and distributed trace-memory networks^45,46^. Because correlation structure can be influenced by global state fluctuations and movement, we interpret these networks as proxies for functional coupling and focus on within-animal, epoch-to-epoch reorganization and region-specific patterns rather than absolute correlation magnitude.

Beyond the immediate application to fear conditioning, COMET enables systematic tests of how distributed cortical networks rapidly reorganize during learning—an empirical gap that remains between influential computational frameworks (for example, Rescorla–Wagner and reinforcement learning models^47,48^) and neurophysiological measurements that capture how network architecture changes in space and time. The ability to map trial-by-trial, region-resolved network transitions at single-cell resolution in freely moving animals provides a route to identify early network signatures of learning, distinguish sensory/motor components from mnemonic components, and constrain mechanistic models of prediction and credit assignment across cortex.

COMET is designed as an open and extensible platform. By releasing head-stage designs, control electronics and analysis software, we aim to facilitate adoption and extension to additional complex behavioral paradigms and animal models (**Supplementary Table 1**), as well as to experimental contexts involving pathological or physiological perturbations that impact distributed cortical networks, such as inflammation, tumor growth, or infection. By enabling longitudinal, cortex-wide recordings in freely moving mice, COMET provides a framework to link spatially localized perturbations to their global functional consequences across the cortex. Several engineering directions could further broaden the accessible experimental space, including weight reduction through alternative structural materials, more compact optical routing to decrease core mass, dual-color capability for concurrent readouts (for example, calcium alongside hemodynamics, neuromodulators, or inflammatory signals), and higher frame rates enabled by next-generation sensors. Finally, integrating optogenetic stimulation would enable causal perturbations during cortex-wide recording, allowing direct tests of whether early network transitions during complex behavior predict — or drive — subsequent behavioral expression.

## Methods

### COMET design, manufacturing and assembly

#### Optics optimization

In applications requiring a large FOV, the selection of the optical system structure is particularly critical. Among these, symmetric systems are widely used due to their unique advantages. Such systems originate from fully symmetric optical systems, which are characterized by structural parameters that are completely symmetric with respect to the aperture stop. The defining feature of a fully symmetric optical system is that if the stop surface is considered as a plane mirror, the two halves of the system on either side of the stop exhibit a mirror-image relationship. When imaging at a certain magnification, the axial aberrations of a fully symmetric system are the sum of the aberrations of the two halves, while the transverse aberrations cancel each other out, resulting in zero net transverse aberration.

For the system under consideration, a fully symmetric optical system is not suitable because asymmetry leads to an imbalance in aberrations. To address this, we employ a system with asymmetric structural parameters to compensate for the changes in aberrations. Although the structural parameters are no longer symmetric, the system retains a symmetric shape in its structure and is thus referred to as a symmetric-type system. This type of system inherits some of the fundamental aberration characteristics of fully symmetric systems, such as the opposite signs of transverse aberrations for the objective and tube lenses on either side of the stop, which results in cancellation, while the axial aberrations have the same sign and thus add up. If the symmetry plane or symmetric parameters are made asymmetric, i.e., increasing one parameter while decreasing the other, the transverse aberrations can be altered without affecting or minimally affecting the axial aberrations. This type of system has a wide range of applications and is particularly suitable for large field of view scenarios.

In the design of this system, particular attention has been paid to match the optical powers of the objective and tube lenses. The optical powers of the objective and tube lenses are essentially equal, and a separated doublet design is used to correct for spherical aberration and longitudinal chromatic aberration. At the same time, the characteristics of the symmetric-type system are utilized to effectively correct transverse aberrations such as coma, distortion, and lateral chromatic aberration. Additionally, by carefully selecting the position of the stop, astigmatism is effectively controlled. To address the field curvature introduced by the large field of view, a negative power field lens is placed near the object plane. This design not only effectively corrects field curvature but also further enhances the imaging quality of the system. In this way, all seven primary aberrations are corrected using just five spherical lenses.

The optical system of the COMET was designed using Zemax OpticsStudio. The goal of the optical system is to produce a miniature microscope with up to 10 mm FOV and single-cell resolution, in which the desired NA is set to be 0.083 in the objective space with minimal field curvature. Three achromatic lenses are designed as the objective module. Two achromatic lenses form the tube lens module. The EWL is placed between for electronic focus adjustment.

#### Filtering optimization

Even excitation across the FOV is ensured by adopting 8 LEDs filtered with 16 excitation filters, each with a size 2.1 × 1.5 ×0.5 mm. Two filters stacked with the non-coating side in contact are placed in front of each LED to achieve a better filtering performance. Instead of using an interference filter, an absorption filters are placed in front of the CMOS sensor. As absorption filters (10 × 10 × 0.1 mm, Kodak Wratten Filters, #54-467, Edmund Optics) are angle-independent, they perform better in the case of diffracted light with larger angles to achieve a better contrast.

#### Illumination optimization

The baseplate module accommodates eight LEDs and sixteen excitation filters (ET470/40x, Chroma). Each LED is paired with two excitation filters stacked with their non-coated surfaces facing each other, together with a paper diffuser, to further narrow the excitation bandwidth and improve illumination uniformity.

#### PCB board design

The circuit schematic and rigid-flex PCB layout are designed using KiCad (version: 9.0.3), a free software suite for electronic design automation. The PCB is divided up into 2 rigid and 1 soft subcircuits. One includes an EWL tuning, head orientation circuit, and a power-over-coax and serializer circuit, the second contains a CMOS image sensor circuit, and the third soft excitation LED circuit contains 8 LEDs to generate an even illumination in the object space across a up to 10-mm field of view. The three subcircuits are connected by a double-sided embedded flex printed circuit.

#### Main body fabrication

The COMET module bodies (objective module body and emission module body) are machined with Magnesium-Aluminum Alloy (1.77 g/cm3, AZ31B) with electrochemical blackening. The sensor module body, baseplate, and protective cap are designed in Autodesk Fusion 360 (educational license) and printed with black resin (FLGPBK04, Formlabs) by a tabletop stereo lithographic 3D printer (Form 3, Formlabs) to produce a lightweight, compact, and low-cost assembly.

#### Assembly

The assembled objective module is attached to the emission module body with three Hex Socket Screw (M1.6×3). The emission module, sensor module, cover, and rigid-flex PCB are fastened together with four M1thread-forming screws (96817a704, McMaster-Carr). The COMET is attached to the baseplate with a M2 set Screw (**Extended Data Fig. 2**).

#### Wiring

A single flexible coaxial cable (50 ohms; CW2040-3650SR, CoonerWire) is used for power, communication, and image data transmission, relying on a passive power-over-coax filter and a serializer/de-serializer pair (TI DS90UB913A/DS90UB914A) for combining and separating dc power, low-speed bidirectional communication, and high-speed unidirectional imaging data. For coaxial cable lengths longer than 2.5 m, an external 6-V power supply should be connected to the UCLA Miniscope DAQ, replacing the power supplied by the USB connection to the COMET. With a total cable diameter down to 0.3 mm (100065-0023, Molex) and compatibility with active and passive, low-torque commutators, this design minimizes the impact of cabling on animal behavior.

#### Data acquisition

The DAQ hardware and software were adapted from the UCLA Miniscope project (http://miniscope.org/index.php/Main_Page), with upgraded firmware (https://github.com/Aharoni-Lab/Miniscope-DAQ-Cypress-firmware) and software (https://github.com/Aharoni-Lab/Miniscope-DAQ-QT-Software) to support video streaming and COMET control. The software provides: excitation LED brightness adjustment, focus control via EWL (electrically tunable lens), real-time ΔF/F calculation and fluorescence trace visualization, configurable frame rate and gain settings, integrated DeepLabCut-Live for real-time pose estimation (via embedded Python). Additionally, we implemented real-time zoom functionality during live monitoring to verify single-neuron activity and ensure proper COMET alignment across the entire FOV.

Future updated versions of all ordering numbers of key components, 3D models and drawings of the customized components, building instructions and video tutorials, and customized recording software, can be found here: https://github.com/ChangliangGuo2/COMET/.

### Animals

#### Mice chosen

Adult C57BL/6J mice (male or female without randomization or blinding) at 8–16 postnatal weeks were applied in the study and were housed in an animal facility (24 °C, 50% humidity) under a 12:12 reversed light-dark cycle. All animal experiments were performed following institutional and ethical guidelines for animal welfare and have been approved by the Institutional Animal Care and Use Committees at Tsinghua University and Peking University. For blood vessel imaging in freely moving mice, C57BL/6J wild-type mice were intravenously injected with FITC–dextran (70 kDa; Sigma) at 25 mg mL⁻¹ in saline, corresponding to a dose of 200 mg kg⁻¹. For functional imaging of neural activity in comparison with the tabletop one-photon widefield system, two-photon microscope imaging, as well as behavioral studies of social interactions and fear conditioning, we used transgenic mice hybridized between Rasgrf2-2A-dCre mice (Jax:022864) and Ai148 (TIT2L-GC6f-ICL-tTA2)-D mice (Jax:030328) expressing Cre-dependent GCaMP6f genetically encoded calcium indicator (GECI). Trimethoprim (TMP; 50 mg kg^−1^ body weight) was injected intraperitoneally ∼1 week before surgery to induce the expression of GCaMP6f in layer 2/3 cortical neurons across the whole brain.

#### Surgical implantation

For implanting a glass imaging window, mice were initially anesthetized with 3.0% (by volume in oxygen) isoflurane in an acrylic box for 3∼5 min and then were head-fixed on a stereotaxic apparatus (Kopf Instruments). Mice were maintained anesthesia with 1.5% isoflurane during the whole surgery period. After removing the scalp and skull piece, a Trapezoid-shaped (a top base of 7 mm, a bottom base of 10 mm, and a height of 7.5-8.0 mm) coverslip was implanted on the craniotomy region, and a titanium headpost was then cemented to the remaining skull over cerebellum for head fixation. Flunixin meglumine (Sichuan Dingjian Animal Medicine Co., Ltd) was applied for perioperative analgesia and postoperative pain care (1.25 mg/kg, subcutaneous injection, one dose daily for three consecutive days). After inflammation was gone and the cranial window became clear (∼2 weeks after surgery), mice were ready for imaging^51^. For implantation of COMET baseplate module, mice with glass imaging window were head-fixed on a customized headpost holder. Real-time Ca^2+^ imaging using COMET system were conducted to find the best focus for the cortex-wide neurons. After that, the baseplate was cemented on the top of imaging window with focus length being fixed (**Extended Data Fig.2**).

### Benchtop validation of optical imaging performance of the COMET

Validation of single-cell performance of COMET with a commercial 2P microscope as a gold standard uses two methods. The first one is aligning the footprints of the neurons from the COMET to the footprints of the neurons from 2P microscope to check the percentage of neurons from the COMET that can be validated in the neurons from 2P microscope. The second method refers to a statistical analysis in terms of df/f and event rate from the COMET and 2P microscope to check whether they follow the same results. The same cortical regions of a head-fixed mouse was imaged with both COMET and 2p microscope. 2P imaging was taken by scanning from 20 µm to 170 µm with a 15 µm step size, with a total of 11 layers captured, denoised with SN2N algorithm. The extracted neurons of 11 layers were taken into account.

### Freely moving animal experiments

#### Trace fear conditioning(TFC)

The trace fear conditioning arena was a chamber (W, 18 cm; D, 18 cm; and H, 30 cm) and a grid floor that consisted of 16 stainless steel rods. A camera (ACT-100A, Coulbourn Instruments) was placed overhead for behavior recording. During the conditioning experiment, the chosen mouse mounted with the COMET was placed in a fear conditioning chamber for a 4-minute free exploration (SPON). Then, the mouse received periodic tone cues (lasting 20 s), tone interval (lasting 18 s), electrical foot shocks (duration 2 s, 0.75 mA), and post-shock interval (lasting 60 s), 5 trials in total. The paradigm ended with a light signal.

#### Social behaviour experiments

In our experimental paradigm, a single mouse equipped with the COMET imaging system was first placed in an open-field arena and allowed to explore for 5 minutes. A second mouse was then introduced, and their interactions were recorded for an additional 5 minutes. The social period was manually annotated with the behavior recording from the behavior camera placed overhead.

#### Open-field test

To test the effects of carrying COMET on freely exploration behavior, four mice were individually introduced into an open field (40 × 40 × 40 cm, length-width-height) for two consecutive sessions (10 min per session). In one session, mice were with only the Baseplate. And in another session, mice were with the Baseplate and COMET. To minimize potential order effects, two mice were initially tested without COMET followed by with COMET, while the remaining two mice received the reverse sequence.

### Data analysis

#### Behaviour video analysis

Videos of mouse behaviour were captured in AVI format using an overhead camera. For the trace fear conditioning paradigm, the mice were tracked using DeepLabCut. Tracking output of running speed (head-movement derived) from each trial was manually verified to ensure tracking accuracy. For the social interaction experiment, the social interaction period was manually scored.

#### Time-lapse neuronal data processing

Captured raw data from the COMET was first removed of salty noise from the dead pixels of the CMOS, and then motion-corrected using the open-source NormCorre algorithm in non-rigid mode. Then the data was denoised with SN2N denoising algorithm, followed by CaImAn pipeline to extract footprints of neurons and calcium activity. The Allen Brain Atlas was used as an anatomical reference for data analysis in this study (http://www.brain-map.org).

#### Matching cells across sessions

Contours computed from CaImAn were normalized by their maximum value and then thresholded at 0.5 to generate compact contours for all cells. Each session was manually aligned using the contours and vasculature simultaneously with custom MATLAB scripts. In addition to linear translations, it was sometimes necessary to radially scale these data as well as to account for slight differences induced by camera placement and different focal positions. For all mice, we aligned and matched three consecutive recording sessions (24 hours apart) where running performance was stable. Contour alignment was performed using CellReg to register cells across all sessions of interest based on their centroid distances and spatial contour correlations. Signal scattering through issue and background noise inherent to single-photon miniature microscopy increases the difficulty of tracking cells across sessions; however, CellReg provides metrics for understanding the quality of identification based on multiple measures, allowing researchers to quantify their confidence in the registration. Distributions of centroid distances and spatial correlation for all cell pairs within 24 μm were computed, yielding a 2D distribution that can be modeled and given a probability threshold to match cell pairs. Cell pairs with a probability of >0.5 for both the centroid and correlation distributions were matched (typically centroid distances of <6 μm and spatial correlation of >0.8), which is expected when recording the same cell.

#### PSF blurring

As a comparison, non-single-cell resolution ROI analysis requires recording the cortex with an optical system that could not achieve single-cell resolution. Instead of developing a real device, we adopted a 50 µm Point spread function to convolve the raw videos captured by COMET. Then, the non-single-cell resolution ROIs are extracted according to the footprints of the neurons from COMET for further comparison analysis.

#### Neural network analysis

Mean degree centrality was computed to assess the relative importance of nodes within the network. Degree centrality, defined as the number of edges connected to a node, was calculated for each node, and the mean value across all nodes was derived to summarize overall network connectivity. This measure provides insight into the network’s global structure, where higher values indicate greater average connectedness among nodes. An undirected weighted network is represented by a symmetric adjacency matrix (*W*), calculated from the Pearson correlation coefficient between all pairs of neurons followed by keeping only correlations with absolute values > 0. Each entry *W*(*i,j*) = *W*(*j,i*) represents the weight of the edge between nodes *i* and *j*. Diagonal elements *W(i,i)* are typically 0 (no self-loops unless specified). The weighted degree centrality (or node strength) of a node is the sum of the weights of all its connected edges. For node *i*, its strength is calculated as: strength*_i_* = ∑*_j_W*(*i, j*). Since the network is undirected, summing either rows or columns of gives the same result. The average weighted degree centrality is simply the mean of all node strengths: 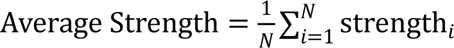, where N is the number of nodes in the network. All analyses were performed using Matlab 2022a.

#### Structure similarity (SSIM)

SSIM is a widely used metric for the assessment of visual quality of images and remote sensing data. We used the ‘ssim()’ function in MATLAB to calculate the similarity between reconstructed images and ground-truth images (obtained by the average of the images from the whole recording after motion correction).

#### Peak Signal-to-Noise Ratio (PSNR)

PSNR is an engineering term for the ratio between the maximum possible power of a signal and the power of corrupting noise that affects the fidelity of its representation. We used the ‘psnr()’ function in MATLAB to calculate the similarity between reconstructed images and ground-truth images (obtained by the average of the images from the whole recording after motion correction).

## Acknowledgements

This work was supported by the National Natural Science Foundation of China (grant no. T2288102 to C.L.), the National Key Research and Development Program of China (grant no. 2023YFC3402600 to Z.D., grant no. 2022YFC3400600 to L.C.), the National Natural Science Foundation of China (grant no. 32571687 to C.G., 32227802 and 82525105 to L.C., grant no. 32422052 and 62305083 to W.Z.), State Key Laboratory of Advanced Manufacturing for Optical Systems (grant no. KLMSKF202402 to C.G.), the Scientific Research Innovation Capability Support Project for Young Faculty (grant no. ZYGXQNJSKYCXNLZCXM-H7 to H. L.), the Young Elite Scientists Sponsorship Program by China Association for Science and Technology (grant no. 2023QNRC001 to W. Z.), and the STI2030-Major Projects (grant no. 2021ZD0203600 to Z.G.).

## Author contributions

C.G. initiated and conceived the research. L.C., C.G., W.Z., Z.G., Q.F., and W.L. supervised the project. Q.F. and C.G. designed, fabricated, and assembled the optical module. C.G., M.X., and G.Q. developed the excitation module. C. G. and M.X. developed the PCB circuit. G.Q. and Y.L. constructed the image processing pipeline. J.L. designed the experiments. J.L., M.X, G.Q., and C.G. performed the experiments and collected the data. C.G., M.X., G.Q., J.L., J.C., Y.L., and Y.H. analyzed the data and prepared the figures. C.G., M.X., and G.Q. prepared the videos. L.Q., X.Z., B.Z., J.Z., X.L., H.L., J.M., B.Z., X.Y., P.Z., Z.L., Z.Z., W.D., J.W., W.W., S.Z., and H.L. participated in discussions during the development of the manuscript. C.G., L.C., and W.Z. wrote the manuscript with input from all authors. All authors participated in the discussions and data interpretation.

---

**Extended Data Fig. 1.**
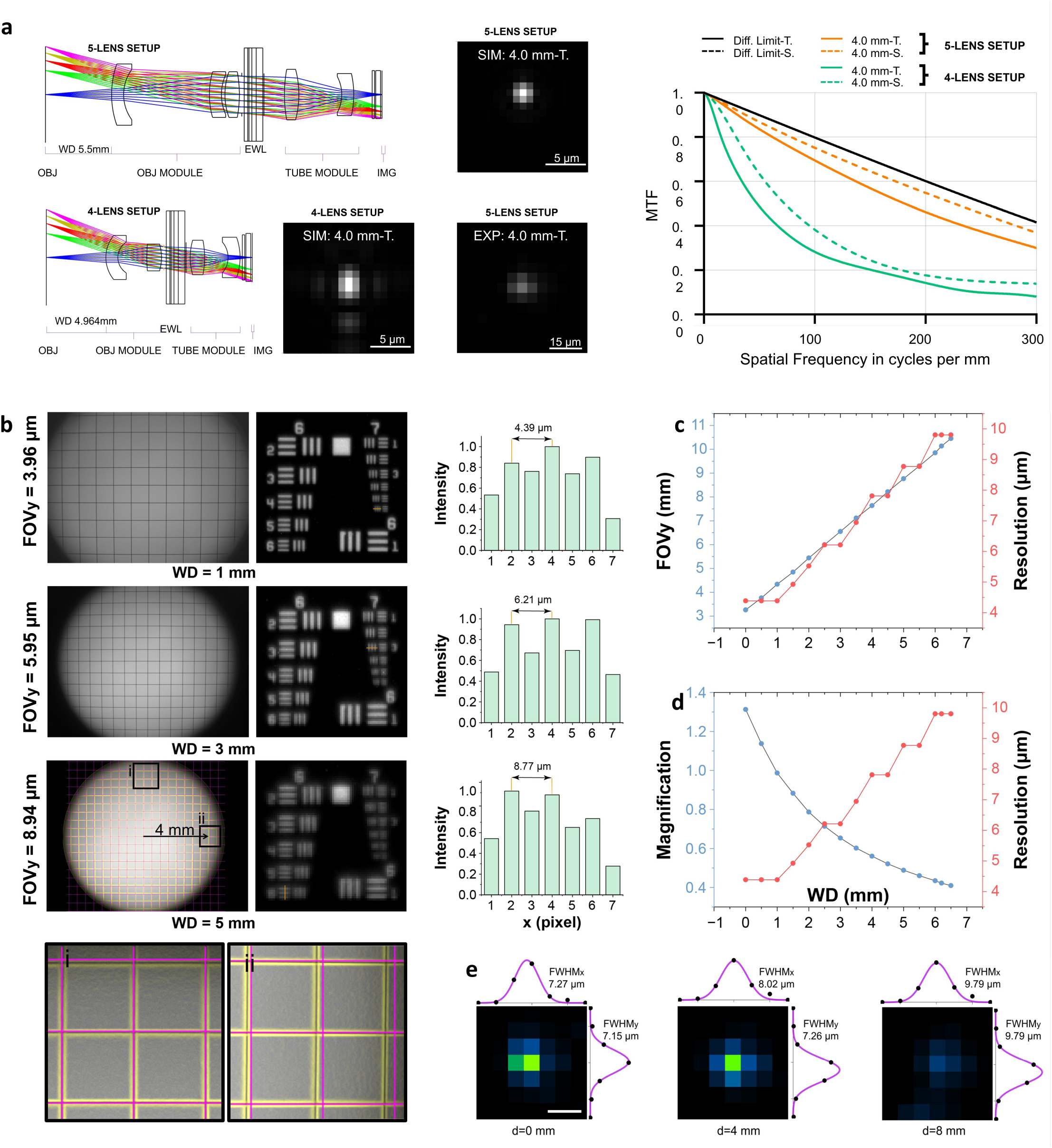
System Characterization: Field of View, Resolution, and Working Distance. **a,** Optical design validation. Zemax simulations comparing 5-lens (27 mm length, 5.5 mm WD, 0.468× mag) and 4-lens (17 mm length) configurations. The 5-lens design maintains near-diffraction-limited performance (PSF/MTF) at 4 mm off-axis, while the compact 4-lens alternative shows degraded resolution. **b,** Field-of-view (FOV) and resolution measurements demonstrated an approximately linear relationship, with 4.35 mm FOV achieving 4.39 µm resolution when the working distance (WD) is 1 mm, 6.55 mm FOV yielding 6.21 μm resolution (WD=3 mm), and 8.77 mm FOV providing 8.77 µm resolution (WD=5 mm). Grid target imaging confirms minimal distortion and field curvature across the 10-mm FOV, with slight edge magnification variation (insets i and ii). **c,** Systematic testing revealed that FOV scaling from 3.26 mm to 10.45 mm resulted in corresponding resolution changes from 4.39 µm to 9.80 μm at the FOV center, as quantified using a USAF resolution target. **d,** Optical magnification decreased inversely with working distance, ranging from 1.3× at 0 mm to 0.45 × at 6.5 mm working distance. **e,** Resolution validation. Single-cell resolution was quantitatively validated by imaging 2-µm fluorescent beads across the entire FOV. The system achieved a full width at half maximum (FWHM) of 7.27 µm (x-axis) and 7.15 µm (y-axis) for a bead at the center (0,0), 8.02 µm (x-axis) and 7.26 µm (y-axis) for a bead 2mm off the center (d=4mm), and 9.79 µm (x-axis) and 9.79 µm (y-axis) for a bead 4mm off the center (d=8mm), consistent with theoretical predictions. Scale bar: 10 µm.

**Extended Data Fig. 2.**
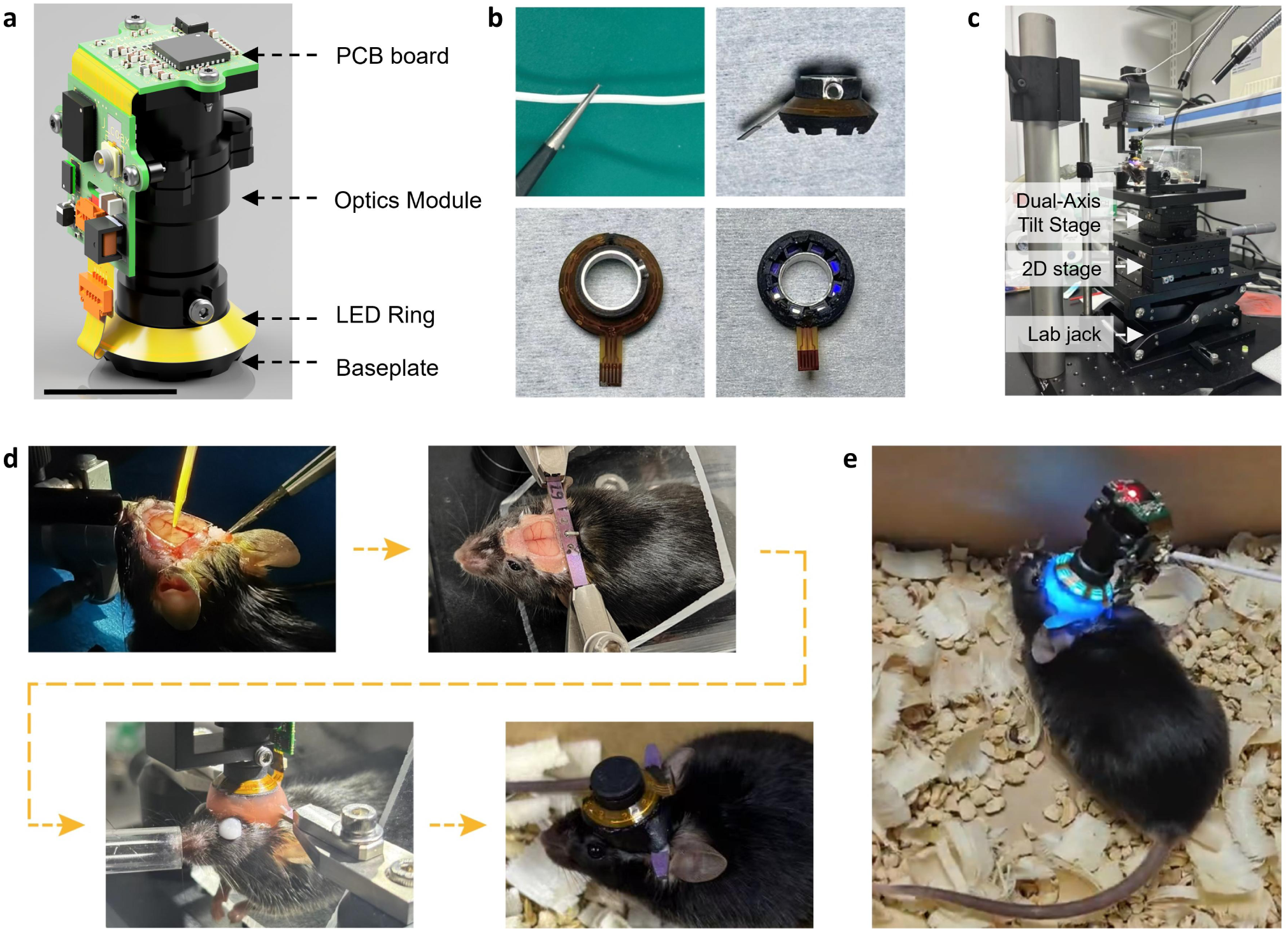
COMET system design, surgical implantation and mounting configuration. **a,** COMET rendering showing compact form factor. **b,** Cabling: 1.3-mm-diameter coaxial cable (50 ohms) with low-torque commutator ensures unrestricted movement; Illumination: Baseplate integrates LED ring with tissue-paper diffusers for uniform excitation across FOV. **c,** Multi-axis stage for fixing the COMET on the skull of the mice. **d,** Cranial window implantation after skull removal, post-surgery fixation, baseplate secured via dental cement, and protective cap shields the window during recovery. **e,** Fully assembled COMET on a freely moving mouse.

**Extended Data Fig. 3.**
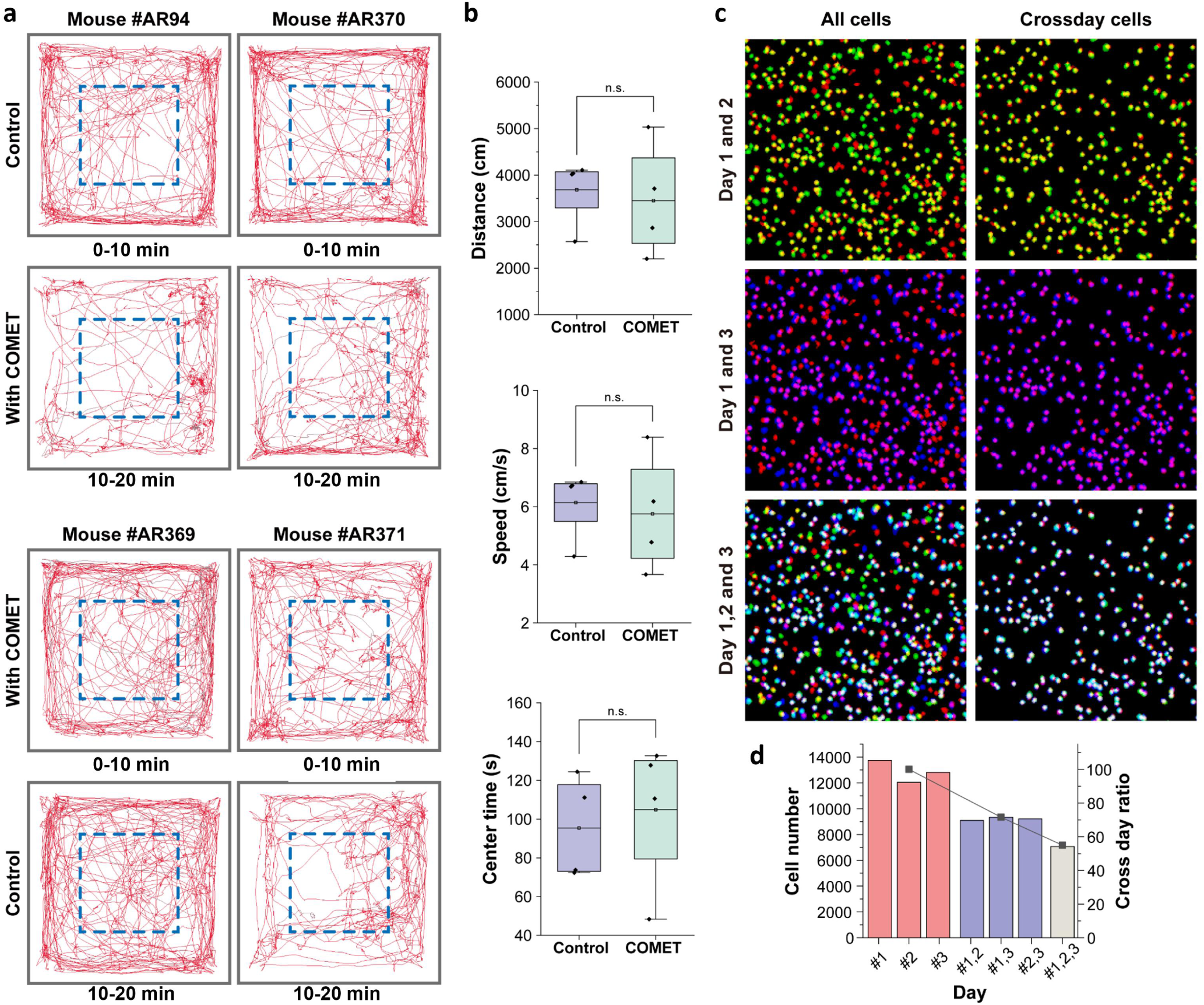
COMET’s lightweight and stable design enable natural behavior and cross-day recordings. **a,** Representative trajectories of 4 mice running for 20 minutes in a 40 × 40 cm^2^ open field, with 2 trials per mouse: 10 minutes for control (no miniscope or cable), and 10 minutes with COMET equipped. Moue #AR94 and #AR360 are tested for control for the first 10 minutes (Row 1) and then 10 minutes with COMET (Row 2), while mouse #AR369 and # AR371 are tested with COMET for the first 10 minutes (Row 3) and then 10 minutes for control (Row 4). Blue dashed boxes show 20 × 20 cm^2^ square as the center area. **b,** Box plot showing total distance traveled (top), median speed (middle), and time spent in the center area (bottom). Colored dots, individual animals. Conditions are compared using Wilcoxon signed-rank test, n.s., p >0.05. **c,** Cross-day stability of COMET imaging. The same cortical cells were tracked for 3 continuous days. Matrix showing overlap of cell footprints in pairs of recordings. Red: cells extracted in day1 recording. Green: cells in day2 recording. Blue: cells in day3 recording. Yellow: overlapping region of day1 and day2. Purple: overlapping region of day2 and day3. White: overlapping region of the three days. **d,** Cell number recorded in the three days, overlapping number and ratio of matched cells of day1-2, day1-3, day2-3, and the three days.

**Extended Data Fig. 4.**
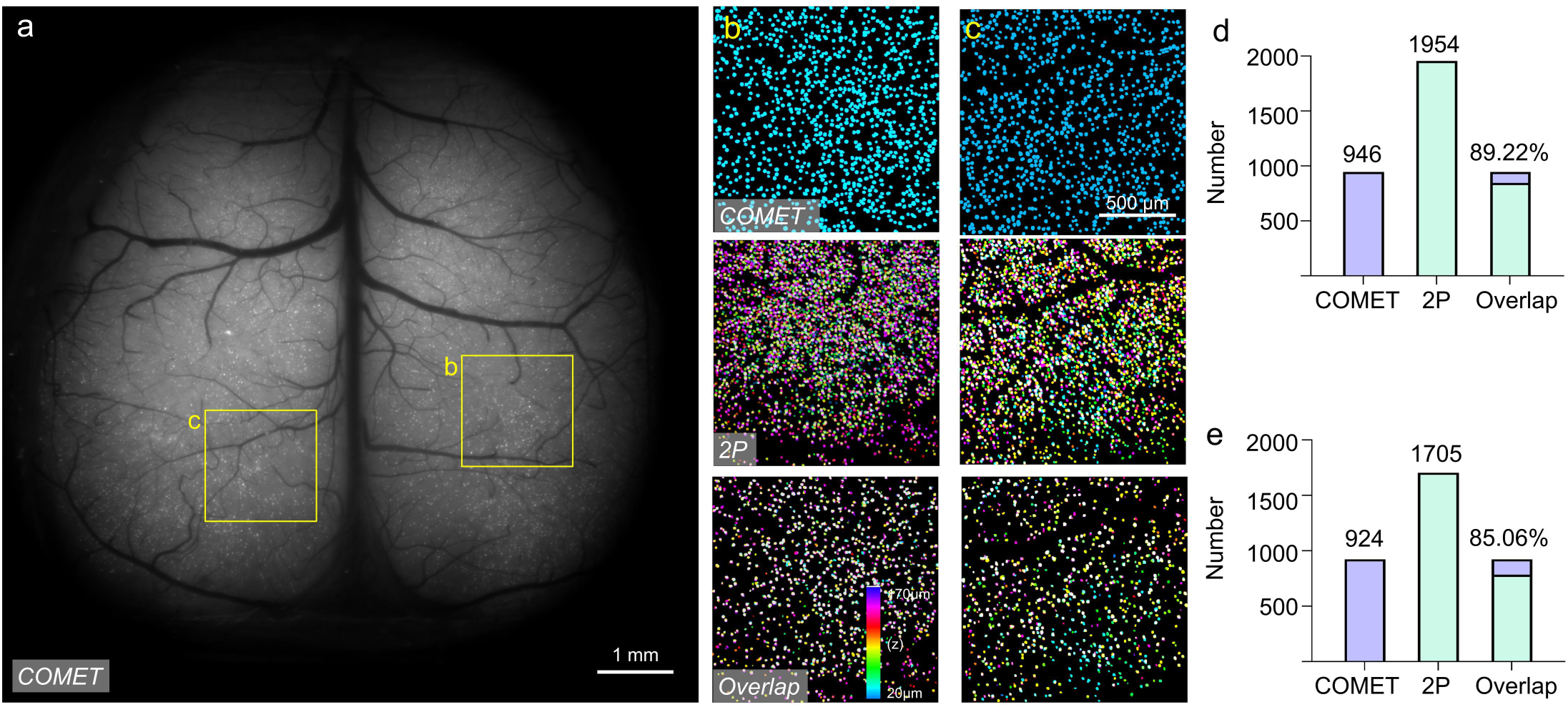
COMET is capable of reliably detecting Ca²⁺ signals from individual neurons across the cortex. **a**, Maximum projection from a 5-min recording from a head-fixed mouse after denoising. Scale bar: 1 mm. **b-c**, Neuron footprints from COMET (top), 11-plane two-photon stack (middle), and overlap (bottom). Scale bar: 500 µm. **d**, 1,954 neurons are identified in two-photon imaging, and 946 neurons were detected in the same regions using COMET. 89.22% of neurons detected by COMET colocalized with those resolved by 2P microscopy. **e**, 1,705 neurons are identified in two-photon imaging, and 924 neurons were detected in the same regions using COMET. 85.06% of neurons detected by COMET colocalized with those resolved by 2P microscopy.

**Extended Data Fig. 5.**
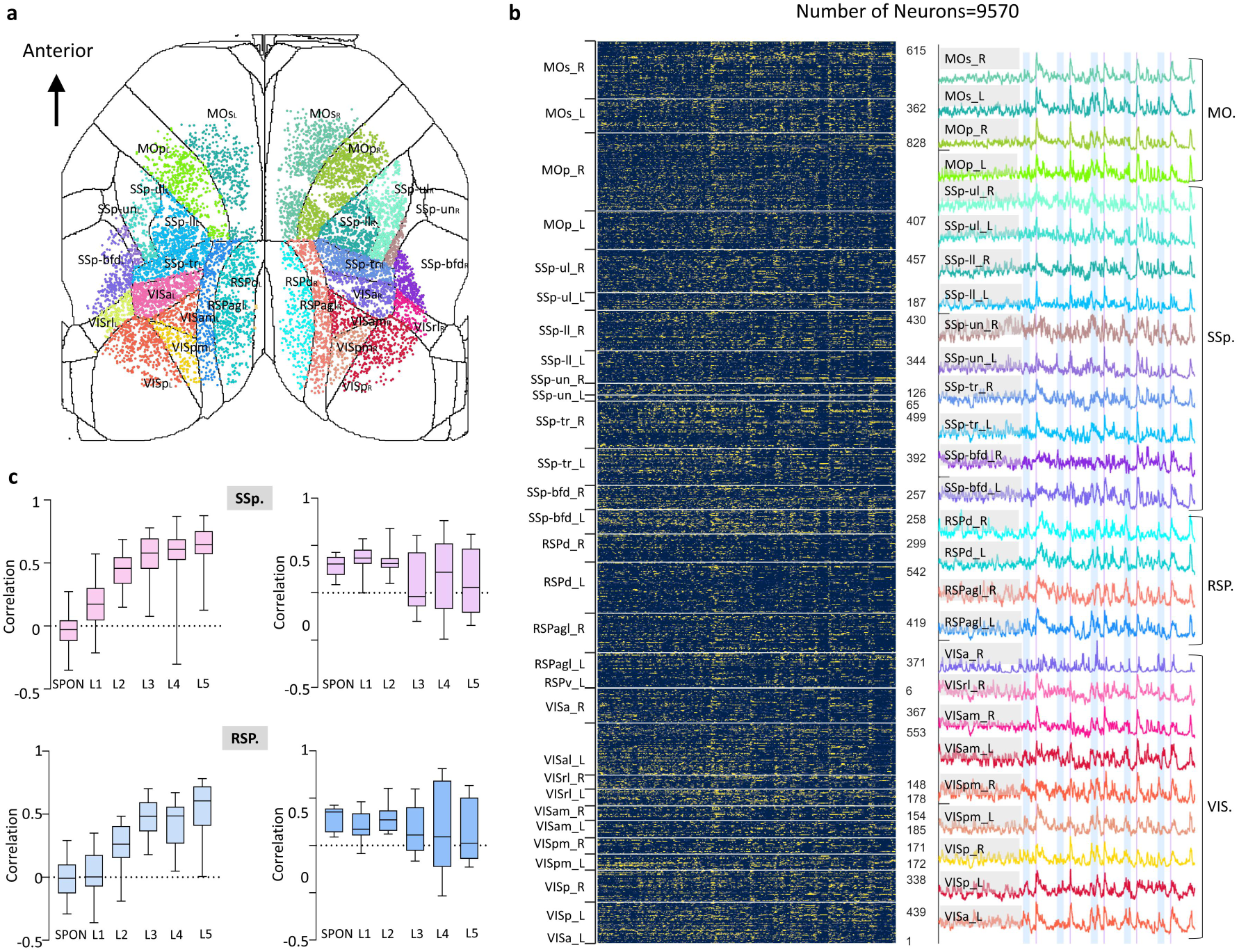
An increase of Pearson correlation between running speed (measured by head movement) and mean neuronal calcium activity during the TFC paradigm. **a,** Spatial distribution of recorded neurons across the FOV. **b,** Normalized calcium activity of 9570 neurons across 30 cortical regions (**left**), and average of normalized calcium activity of the neurons in each cortical region (**right**). Mean of normalized calcium activity by cortical region (grouped into 4 functional domains: MO [motor], SSp [somatosensory], RSP [retrosplenial], VIS [visual]). **c,** Running-activity correlation analysis. Single-cell resolution Pearson correlation (r) during spontaneous (SPON) and post-shock (L1-L5) periods (**left**) and ROI-based correlation (non-single-cell) for comparison (**right**) in SSP. And RSP. Cortical regions.

## Supplementary Information for

**Supplementary Video 1.**
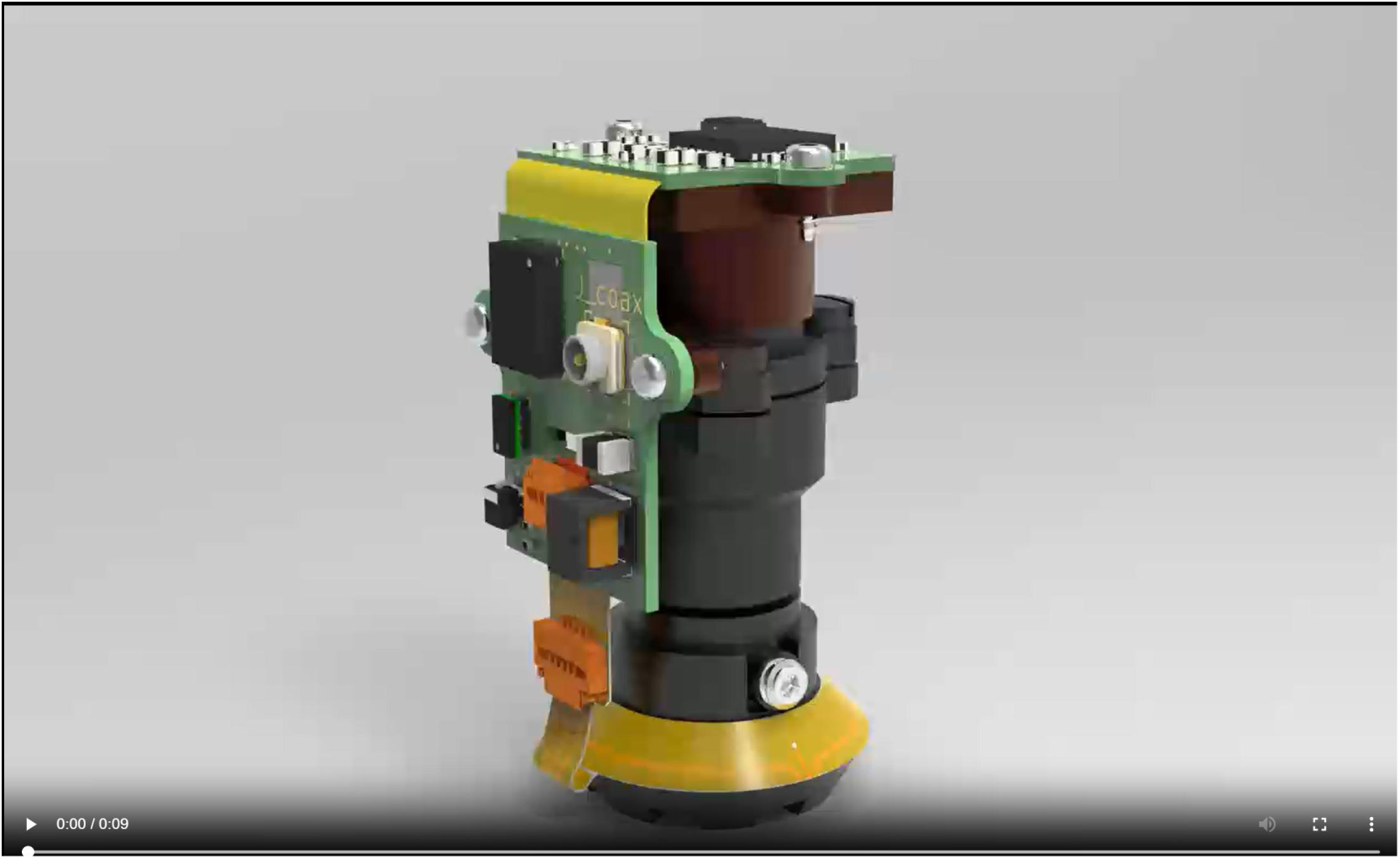
Exploded view of COMET, related to Fig. 1.

**Supplementary Video 2.**
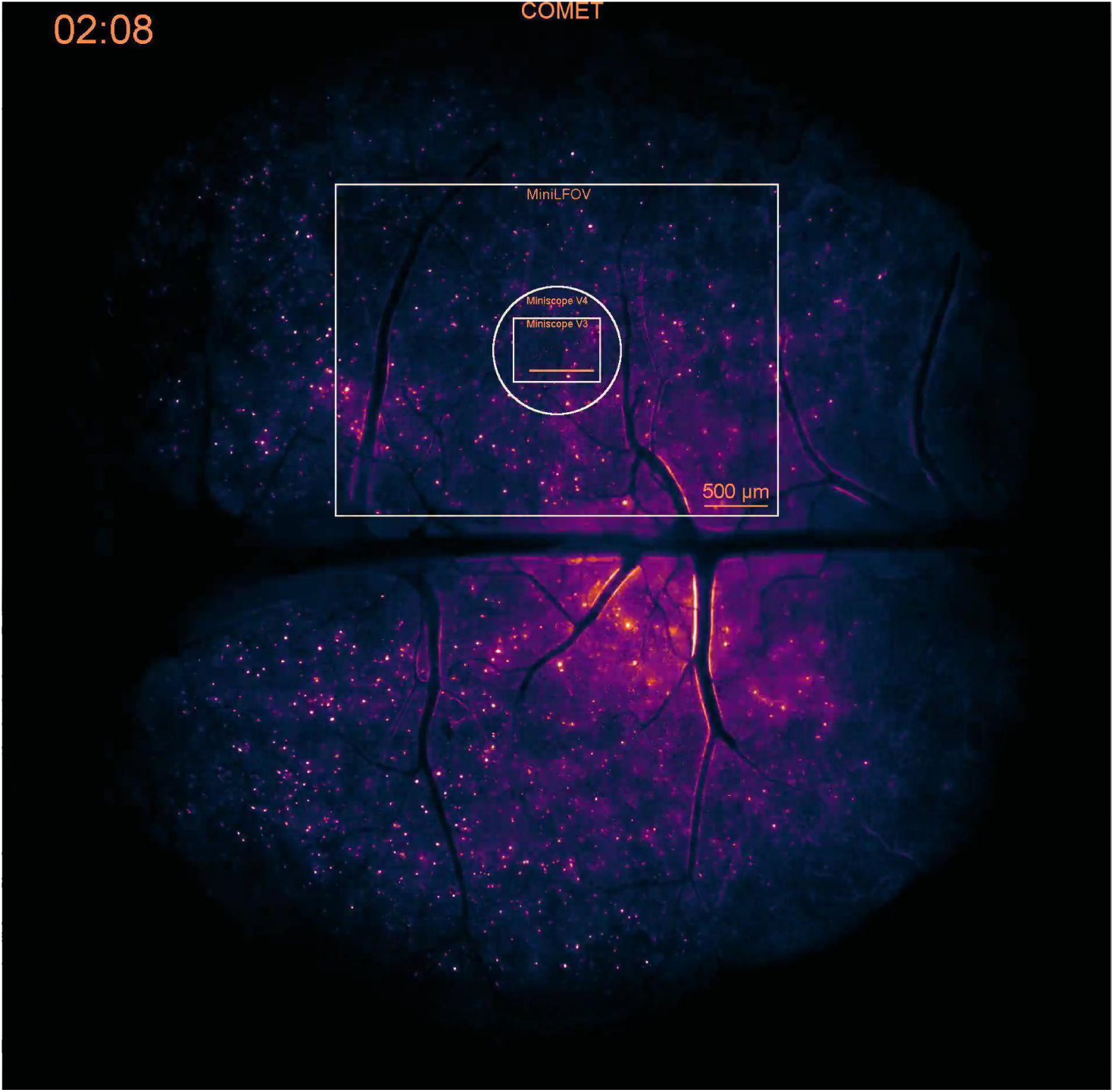
COMET enables whole-cortical imaging with single-cell resolution, related to Fig. 2.

**Supplementary Video 3.**
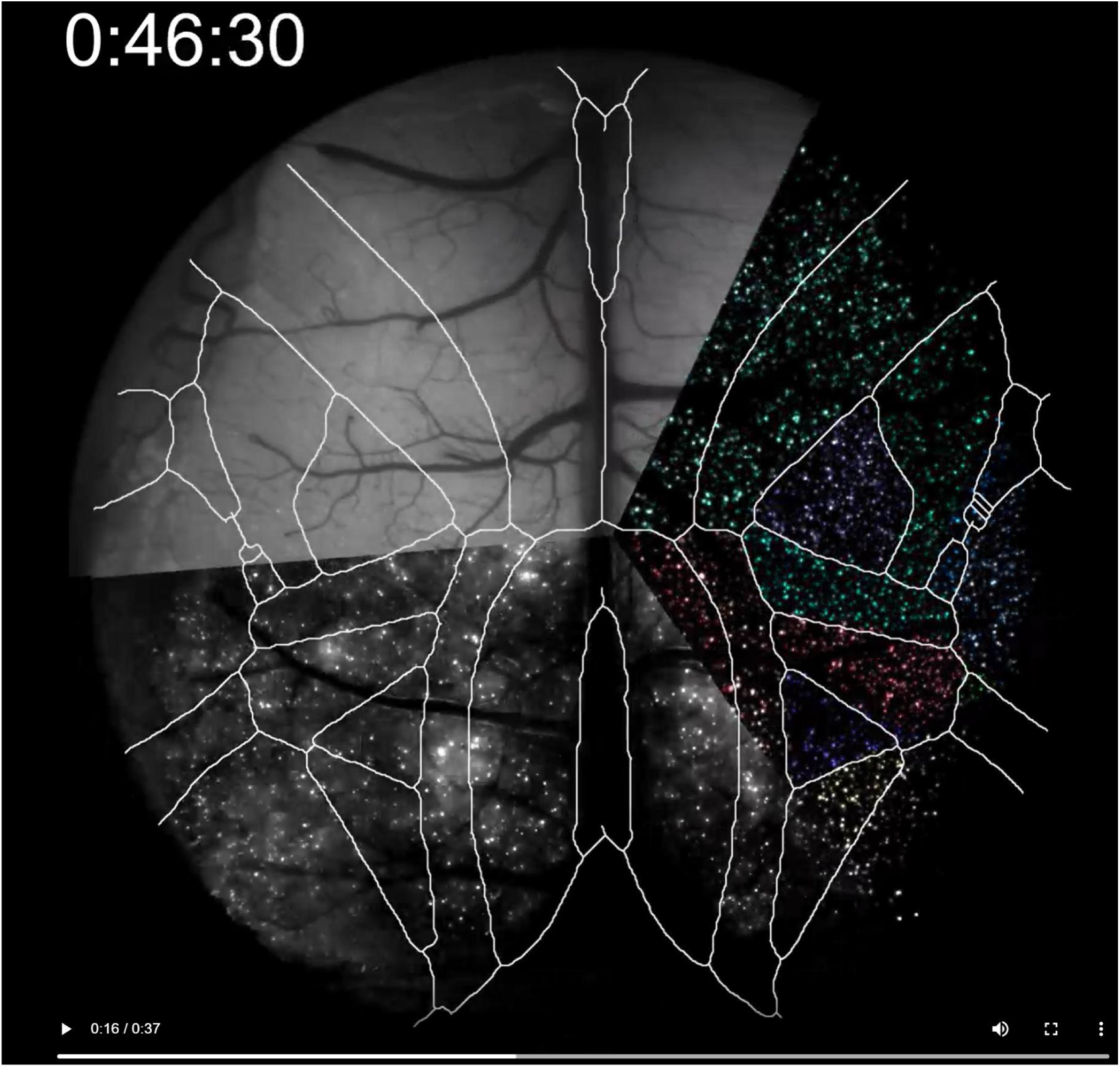
COMET’s computational pipeline enables high-fidelity neuron extraction, related to Fig. 2.

**Supplementary Video 4.**
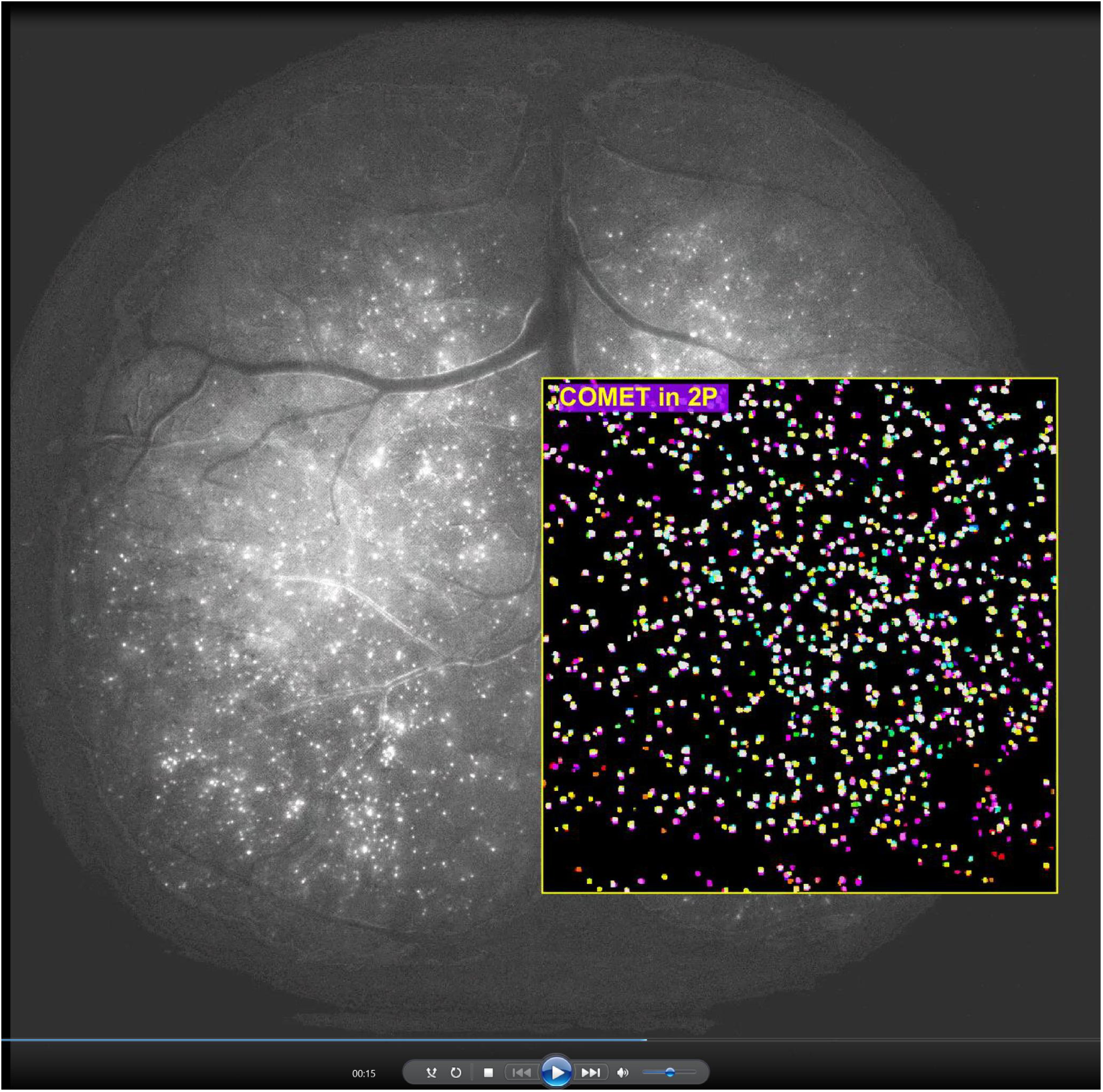
COMET is validated with bench-top 2P microscope, related to Fig. 2.

**Supplementary Video 5.**
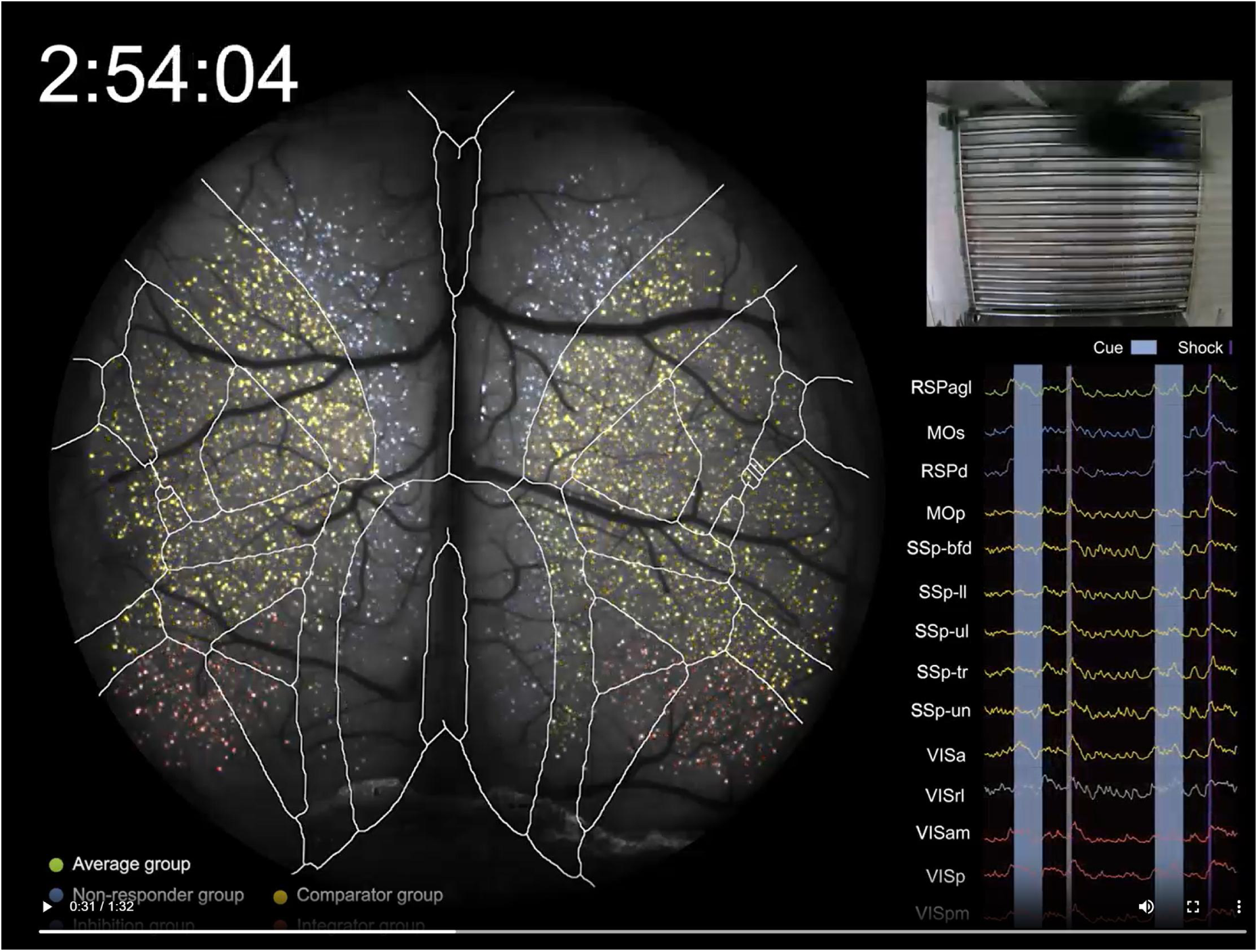
Functional heterogeneity in cortical activity during TFC, related to Fig. 4.

**Supplementary Table 1.**
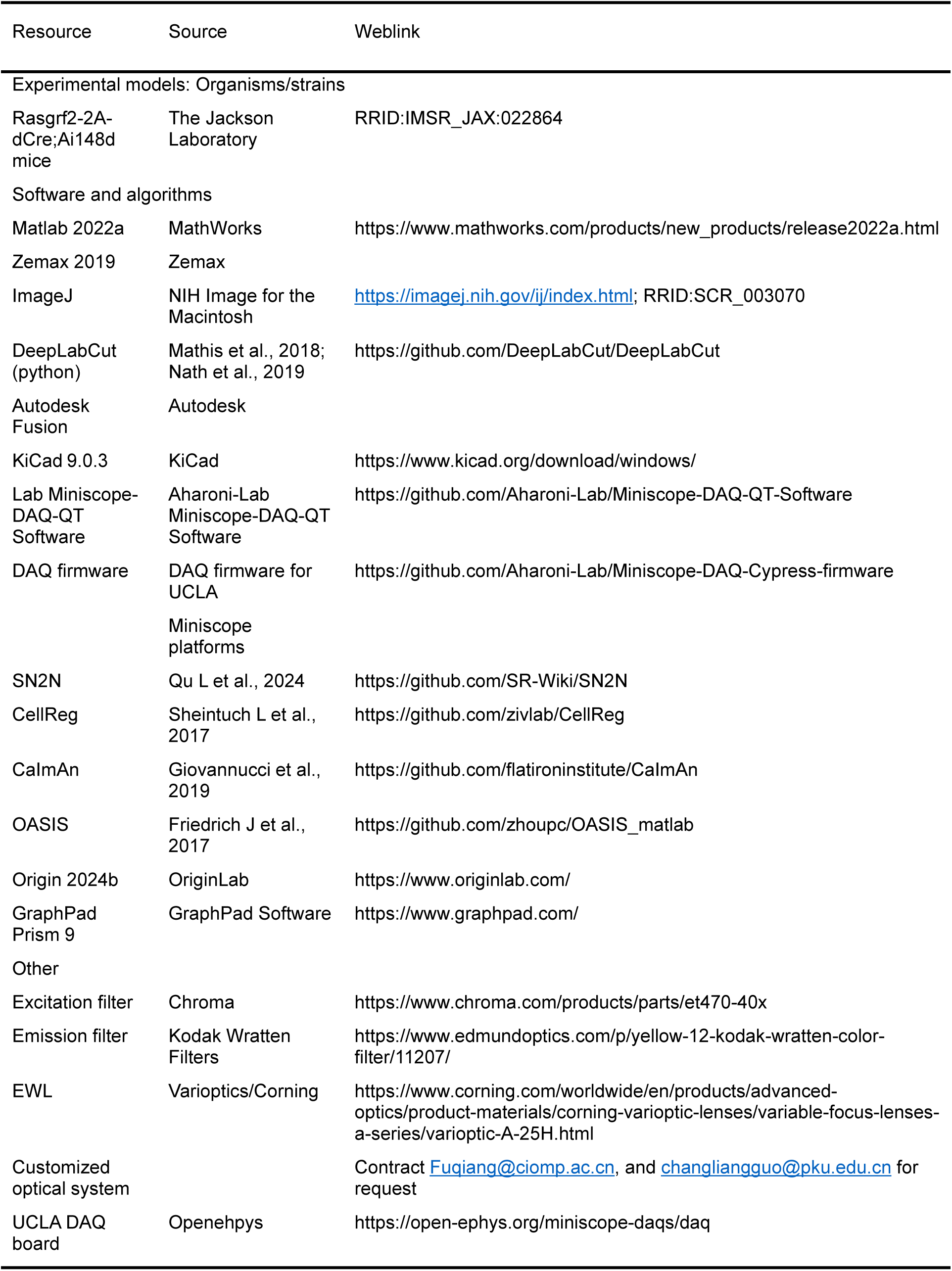
List of resource for COMET.

